# Age-related MICOS Complex Dysregulation Impairs Mitochondrial 3D Architecture and Metabolic Homeostasis in the Liver

**DOI:** 10.1101/2024.06.20.599846

**Authors:** Sepiso K. Masenga, Alexandria Murphy, Prasanna Venkhatesh, Zer Vue, Ashlesha Kadam, Andrea G. Marshall, Benjamin Rodriguez, Estevão Scudese, Brenita Jenkins, Amber Crabtree, Praveena Prasad, Edgar Garza-Lopez, Han Le, Ky’Era V. Actkins, Elma Zaganjor, Nelson Wandira, Jeremiah Afolabi, Prasanna Katti, Chantell Evans, Young Do Koo, Dhanendra Tomar, Mark A. Phillips, David Hubert, Chandravanu Dash, Pooja Jadiya, Olujimi A. Ajijola, Magdalene Ameka, Okwute M. Ochayi, Eric Wang, Quinton Smith, Ronald McMillan, Annet Kirabo, André Kinder, Tyne W. Miller-Fleming, Bret Mobley, Julia D. Berry, Nathan Winn, Vernat Exil, Anita M. Quintana, Kit Neikirk, Jenny Schafer, Sean Schaffer, Oleg Kovtun, Mohd Mabood Khan, Calixto Pablo Hernandez Perez, Margaret Mungai, Melanie R. McReynolds, Antentor Hinton

**Affiliations:** Department of Molecular Physiology and Biophysics, Vanderbilt University, Nashville, TN, 37232, USA; Department of Physiological Sciences, Pathology and Microbiology, HAND research group, Mulungushi University School of Medicine and Health Sciences, Livingstone, Zambia; Department of Medicine, Division of Genetic Medicine, Vanderbilt University Medical Center, Nashville, TN 37232, USA; Department of Biochemistry and Molecular Biology, The Huck Institute of the Life Sciences, Pennsylvania State University, State College, PA 16801; Department of Internal Medicine, Section of Cardiovascular Medicine, Wake Forest University School of Medicine, Winston-Salem, NC 27157 USA; Department of Biomedical Sciences, School of Graduate Studies, Meharry Medical College, Nashville, TN 37208-3501, USA; Department of Biology, Indian Institute of Science Education and Research (IISER) Tirupati, AP, 517619, India; Department of Cell Biology, Duke University School of Medicine, Durham, NC, 27708, USA; Division of Endocrinology, Diabetes and Metabolism, Department of Medicine, David Geffen School of Medicine and UCLA Health, University of California-Los Angeles, Los Angeles, CA, 90095, USA; Department of Integrative Biology, Oregon State University, Corvallis, OR, 97331, USA; The Center for AIDS Health Disparities Research, Meharry Medical College, Nashville, TN; Department of Microbiology, Immunology, and Physiology, Meharry Medical College, Nashville, TN; Department of Biochemistry, Cancer Biology, Pharmacology and Neuroscience, Meharry Medical College, Nashville, TN; UCLA Cardiac Arrhythmia Center, University of California, Los Angeles, CA, USA; Department of Medical Microbiology and Immunology, KAVI-ICR, University of Nairobi, Nairobi, Kenya; Department of Physiology, Faculty of Basic Medical Sciences, Baze University Abuja, Nigeria; Department of Chemical and Biomolecular Engineering, University of California, Irvine, CA, 92697, USA; Artur Sá Earp Neto University Center - UNIFASE-FMP, Petrópolis Medical School, Brazil; Department of Pathology, Vanderbilt University Medical Center, Nashville, TN, 37232, USA; Department of Internal Medicine, Section of Gerontology and Geriatric Medicine, Sticht Center for Healthy Aging and Alzheimer’s Prevention, Wake Forest University School of Medicine, Winston-Salem, NC; Department of Pediatrics, Division of Cardiology, St. Louis University School of Medicine, Missouri, St. Louis, USA; Department of Biological Sciences, Border Biomedical Research Center, The University of Texas at El Paso, El Paso, Texas, 79968, USA; Department of Cell and Developmental Biology, Vanderbilt University, Nashville, TN, 37232, USA; Department of Chemistry, Vanderbilt University, Nashville, TN, 37232, USA; Department of Medicine, Division of Clinical Pharmacology, Vanderbilt University Medical Center, Nashville, TN 37232, USA; Department of Medicine, Division of Cardiology, Vanderbilt University Medical Center, Nashville, TN 37232, USA

**Keywords:** Aging, 3D Structure, Mitochondria, Metabolism, MICOS Complex, Liver Disease, *CHCHD6*, *CHCHD3*, *OPA1*

## Abstract

**Background & Aims:** Aging is associated with a significant decline in mitochondrial function in the liver, leading to an increased risk of liver disease. This study examines age-related changes in the mitochondrial structure of human and murine livers using a combination of Serial Block-Face Scanning Electron Microscopy (SBF-SEM) and mass spectrometry approaches.

**Methods:** This study integrates mitochondrial structure analysis in a murine model with an analysis of liver architecture, lipogenesis, and genetically regulated gene expression in human cohorts. We explored the Mitochondrial Contact Site and Cristae Organizing System (MICOS) complex using SBF-SEM, three-dimensional reconstruction with Amira software, and mass spectrometry techniques.

**Results:** Aging leads to a reduction in mitochondrial size and complexity, resulting in changes in the metabolomic and lipidomic profiles of murine liver cells that are comparable to those observed in aged human samples. We find that genetically modeled expression of MICOS complex genes *OPA1* and *CHCHD3* is associated with chronic liver disease phenotypes within a large biobank population. Furthermore, we observed dysregulated mitochondrial calcium handling and increased oxidative stress due to the disruption of the MICOS complex.

**Conclusion:** Our study highlights the age-associated decline in mitochondrial complexity and metabolic regulation within the aging murine liver and the human population. We have identified that these changes are partially attributable to the age-related loss of the MICOS complex.

**Impact and implications:** This study offers new insights into the changes to mitochondrial ultrastructure that occur during aging. Using SBF-SEM, the quantification of young and aged murine mitochondrial structure was performed, which had previously been an underexplored avenue for measuring mitochondrial changes. The discovery of mitochondrial ultrastructural changes, in conjunction with measurements of age-associated metabolic alterations and gene association data, provides a model for how changes in MICOS expression may modulate age-related impairment of hepatic mitochondria. These results provide a new model by which changes in MICOS protein expression may both cause and be a potential therapeutic target for age-related impairment in hepatic function.

**Highlights:** Decreased modeled expression of *CHCHD3* in individuals of European genetic ancestry is linked to liver transplant and cirrhosis, while decreased modeled expression of *OPA1* in individuals of African genetic ancestry is associated with chronic liver disease and cirrhosis.

Aging alters liver lipid accumulation, MICOS mRNA levels, and disease markers.

Aging reduces the volume and complexity of murine liver ultrastructure.

Aging and diet significantly alter the MICOS complex in mice.

Knockdown of *Mic60* and *Chchd6* lowers Ca^2+^ uptake, retention, and induces oxidative stress in HepG2 cells.

**Graphical Abstract:** 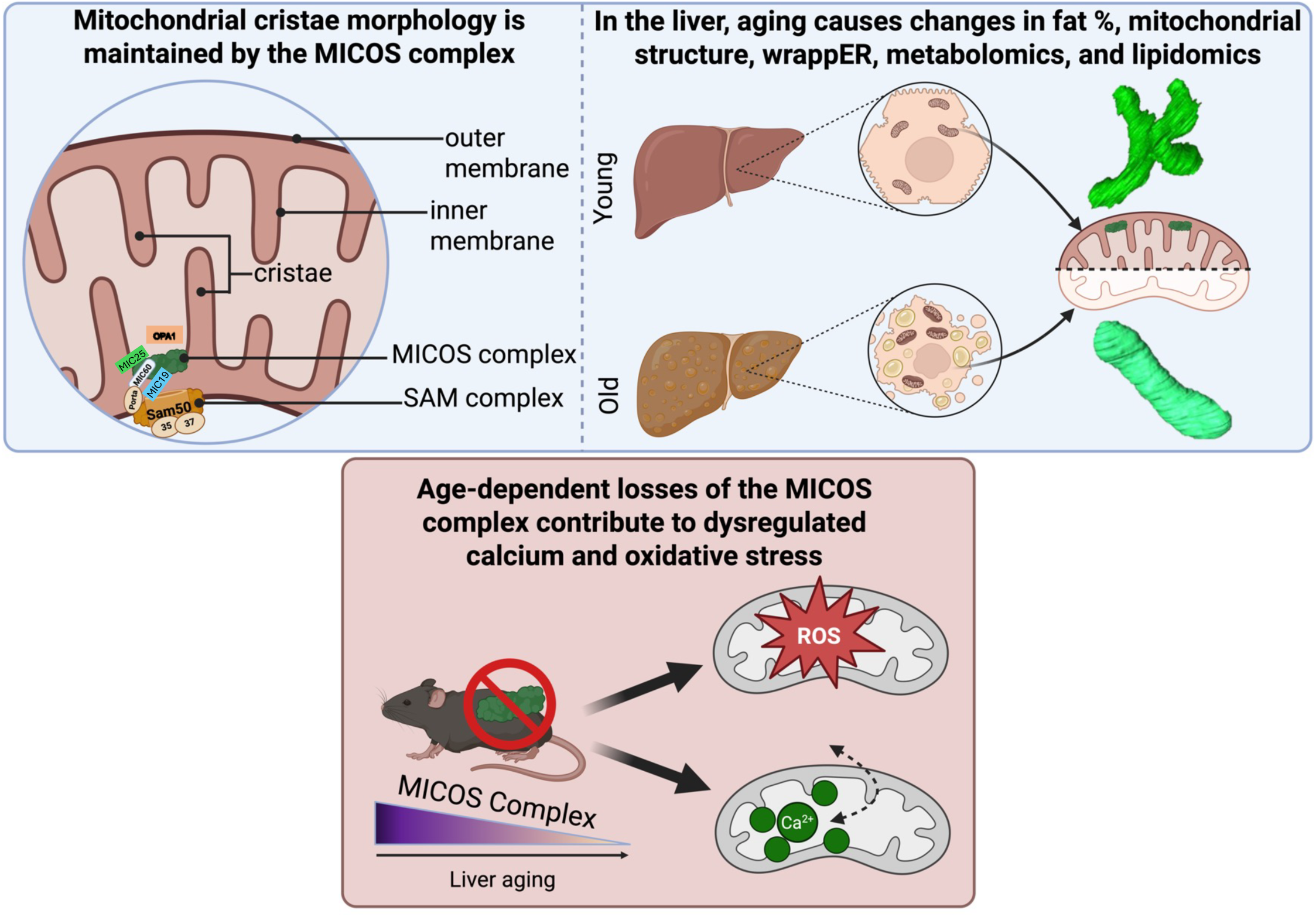

Liver aging causes metabolic, lipidomic, and mitochondrial structural alterations, reflecting age-dependent losses in the MICOS complex. Key components of the MICOS complex (MIC60, CHCHD3 and CHCHD6) are illustrated.

## Introduction

Mitochondria, which are essential for energy production, calcium regulation, and various metabolic pathways, undergo dynamic structural changes through fusion and fission to maintain their function.^1–5^ Aging disrupts these dynamics, impairing mitochondrial ultrastructure and contributing to various diseases.^6–8^ Traditional 2D imaging (e.g. TEM) reveals cristae alterations, but 3D techniques (e.g. SBF-SEM) uncover novel morphologies (e.g., donut-shaped mitochondria, nanotunnels), which remain underexplored in aging tissues.^9–16^ In many organs and experimental models, researchers have not yet performed 3D reconstruction, creating a gap in our understanding of 3D mitochondrial structural changes and their molecular regulators across the aging process in different tissues.

The liver, a metabolic hub critical for detoxification, lipid/glucose homeostasis, and hormone metabolism, experiences age-related declines in mitochondrial function, lipid accumulation, and increased susceptibility to hepatic diseases (e.g., cirrhosis, steatohepatitis, and metabolic dysfunction-associated steatotic liver disease (MASLD), among other hepatic conditions.^17,18^ These pathologies correlate with mitochondrial structural defects, including volume loss, cristae disorganization, and dysregulated dynamics governed by proteins such as MFN1/2, OPA1, and DRP1.^1,19–23^ Additionally, the mitochondria contact site and cristae organizing system (MICOS) complex, which is vital for cristae architecture,^24^ plays a role in age-related dysfunction.^25–28^ Here, we integrate 3D mitochondrial reconstruction, metabolomics, and murine models (juvenile vs. aged) to investigate how aging reshapes hepatic mitochondria. We explore MICOS dysregulation, lipid-driven structural changes, and their consequences on calcium/ROS homeostasis, linking these to transcriptional and metabolic shifts in human aging livers. This work bridges the gaps in understanding 3D mitochondrial remodeling and its role in hepatic aging.

## Materials and methods

### Testing genetically modeled gene expression of MICOS genes with liver disease phenotypes in BioVU

BioVU is a biorepository of matched genotype and de-identified electronic health record (EHR) data curated by the Vanderbilt Institute for Clinical and Translational Research at Vanderbilt University Medical Center. Details regarding the BioVU program, including oversight, patient engagement, and ethical considerations, have been previously discussed.^29,30^ Briefly, this opt-in program collects leftover blood samples from routine patient care visits at clinics across Tennessee, which are then processed for genotyping. BioVU currently houses samples for 344,467 individuals, with ongoing sample collection.

Genotype data for 94,474 individuals were generated on the Illumina Multi-Ethnic Genotyping Array (MEGA^EX^) as previously described.^31^ Quality control procedures included filtering for SNPs and individual call rates, sex discrepancies, and excessive heterozygosity. We used principal component analysis to identify individuals of European and African genetic ancestry, using the 1000 Genomes populations as reference.^32,33^ After imputation using the Michigan Imputation server and the Haplotype Reference Consortium (HRC) reference panel, genotype data underwent additional quality control procedures, including filtering for imputation quality, minor allele frequency, and Hardy-Weinberg Equilibrium.^34,35^ After removing genetically related individuals, we identified 65,363 individuals of European genetic ancestry and 12,313 individuals of African genetic ancestry for analysis. We calculated genetically regulated gene expression (GREX) in BioVU individuals using models built from Genotype-Tissue Expression (GTEx) version 8 project data, which includes genotype and matched transcriptome data for 838 donors across 49 distinct tissues.^36^ The best-performing GREX models based on the highest predictive performance (r^2^) from the JTI (Joint-Tissue Imputation) approach, as implemented in the TIGAR tool, were selected for each gene.^37–39^

To assess the relationship between *CHCHD3, CHCHD6*, and *OPA1* GREX and liver disease, we extracted liver disease case status from the de-identified EHR information of BioVU participants. Specifically, we tested three phenotypes or phecodes mapped from ICD9/10 (International Classification of Diseases, 9th and 10th editions) billing codes: cirrhosis of liver without mention of alcohol (571.51), chronic liver disease and cirrhosis (571), and liver replaced by transplant (573.2). These phenotypes were selected because we were able to identify at least 500 cases within the BioVU population. Previous studies have described the mapping of phecodes to ICD9/10 codes, and researchers can access these mappings through the PheWAS package in R (version 0.99.5-2 and 3.6.0, respectively) or online (https://phewascatalog.org/phewas/#home).^40,41^ We required individuals to have at least two documented instances of the liver disease codes on unique dates within the medical record to be considered a case. We defined controls as individuals with no liver disease codes in their medical records. We used logistic regression analysis to examine the association between *CHCHD3*, *CHCHD6*, and *OPA1* GREX (predictor variables) and the status of liver disease (outcome variable). Covariates included in the regression models were principal components (PC1-10), sex, current age, median age of the medical record, and genotype batch in the ancestry-stratified analyses. We applied the most stringent global Bonferroni p-value by adjusting the p-value for all gene-tissue GREX pairs tested (n=80) and all phenotypes tested (n=3, p=0.05/240). Because the GREX models across tissues are highly correlated, we also generated a less stringent within-tissue Bonferroni p-value adjustment, correcting the p-value for the number of genes tested (n=3) and the number of phenotypes tested (n=3, p=0.05/9). Nominal significance was considered p<0.05.

### Sex As a Biological Variable

In this study, sex was considered as a biological variable in the experimental design. All initial experimental procedures and analyses were performed with attention to potential sex-dependent differences. As TEM analyses revealed minimal sex-dependent differences in the assessed parameters, and given the resource-intensive nature of SBF-SEM, subsequent deep ultrastructural and functional analyses were conducted using male murine cohorts. While sex-specific effects on aging and longevity have been reported and may influence mitochondrial morphology, this study focuses on characterizing aging-associated mitochondrial alterations in a male model.

### Human Cohort

The Vanderbilt University Institutional Review Board (IRB) approved the collection of human liver tissue under the study title “Mitochondria in Aging and Disease - study of archived and autopsy tissue” (IRB No. 231584). Samples came from participants who provided informed consent to participate in this study. We collected samples from participants who died from natural causes or accidents without underlying conditions affecting the liver.

### Animal Care and Maintenance

Per protocols previously described,^43^ the care and maintenance of the male C57BL/6J mice conformed to the National Institute of Health’s guidelines for the use of laboratory animals. We used six male mice for this study: three aged 3 months and three aged 2 years. The University of Iowa’s Institutional Animal Care and Use Committee (IACUC) or the University of Washington IACUC approved the housing and feeding of these mice. We achieved anaesthesia using a mixture of 5% isoflurane and 95% oxygen.

### Oil Red O

We cut OCT blocks into 7 μm thick sections, affixed to glass slides, allowed them to equilibrate to room temperature for 10 minutes, and then stained with Oil Red O (Sigma-Aldrich) as previously described.^44^ We used Student’s t-test to assess statistical significance, ** indicates p< 0.01; and *p< 0.05.

### Bile Acid

From frozen liver tissue, 100 mg of tissue homogenized in 75% ethanol, was incubated for 2 hours, and then centrifuged at 6000g for 10 minutes. Once prepared, we measured bile with the Mouse Total Bile Acids Assay Kit (Crystal Chem) according to the manufacturer’s instructions. We used Student’s t-test to assess statistical significance; ** indicates p < 0.01, and *indicates p < 0.05.

### Triglyceride Levels

We measured the triglyceride levels in the liver and in serum collected after a 6 hour fast using the EnzyChrom™ Triglyceride Assay Kit (BioAssay Systems), with triglyceride extraction performed using a solution of isopropanol and Triton X 100 as described previously.^45,46^ We used Student’s t-test to assess statistical significance; ** indicates p < 0.01, and *indicates p < 0.05.

### Quantification of TEM Micrographs and Parameters Using ImageJ

As per established protocols, we fixed samples in a manner to avoid any bias.^47^ Following preparation, tissue was embedded in 100% Embed 812/Araldite resin with polymerization at 60 °C overnight. After collecting ultrathin sections (90–100 nm), we post-stained them with lead citrate and imaged them using a JEOL 1400+ at 80 kV, equipped with a GatanOrius 832 camera. The National Institutes of Health (NIH) *ImageJ* software was used for quantification of TEM images, as described previously.^14,48^ We used Student’s t-test to assess statistical significance; ** indicates p< 0.01, and *p< 0.05.

### SBF-SEM Processing of Mouse Liver Tissue

We performed SBF-SEM according to previously defined protocols.^10,47,48^ Anesthesia was induced in male mice using 5% isoflurane. After skin and hair removal, the liver was treated with 2% glutaraldehyde in 100 mM phosphate buffer for 30 minutes. It was then dissected into 1-mm³ cubes and further fixed in a solution containing 2.5% glutaraldehyde, 1% paraformaldehyde, and 120 mM sodium cacodylate for 1 hour.

Fixation and subsequent steps were collected onto formvar-coated slot grids (Pella, Redding, CA), stained and imaged as previously described.^10,47,48^ This includes tissue washing with 100 mM cacodylate buffer, incubation in a mixture of 3% potassium ferrocyanide and 2% osmium tetroxide, followed by dehydration in an ascending series of acetone concentrations. We then embedded the tissues in Epoxy Taab 812 hard resin. We performed sectioning and imaging of the sample using a VolumeScope 2 SEM (Thermo Fisher Scientific, Waltham, MA). Conventional TEM analysis was performed on 300–400 serial sections from each sample, following staining and imaging protocols. Subsequently, analysis via imaging was performed under low-vacuum/water-vapor conditions with a starting energy of 3.0 keV and beam current of 0.10 nA. Sections of 50 nm thickness were cut, allowing for imaging at a spatial resolution of 10 nm × 10 nm × 50 nm.

### Segmentation and Quantification of 3D SBF-SEM Images Using Amira

SBF-SEM images were manually segmented in Amira to perform 3D reconstruction, as described previously^10^. We used and analyzed between 300 and 400 slices, and the individual performing the analysis remained blinded to all other details. We collected a total of 250 mitochondria across from each of three mice for each quantification. For 3D reconstruction of liver cells, 10 cells and a total of approximately 200 mitochondria are used, which meets field standards for fulfilling power requirements to determine significant differences between groups. Quantification of 3D structures was performed using the Amira software, which included built-in parameters or previously described measurements.^10^ We performed Mann–Whitney tests for statistical analysis, and significance values indicate **P ≤ 0.01, ****P ≤ 0.001, and ns, not significant.

### LCMS Methods for Metabolomics

Frozen tissues were weighed, ground with a liquid nitrogen in a cryomill (Retsch) at 25 Hz for 45 seconds, before extracting tissues 40:40:20 acetonitrile: methanol: water +0.5% FA +15% NH4HCO3.^49^ with a volume of 40mL solvent per 1mg of tissue, vortexed for 15 seconds, and incubated on dry ice for 10 minutes. We then centrifuged the tissue samples at 16,000 g for 30 minutes. The supernatants were transferred to new Eppendorf tubes and then centrifuged again at 16,000 g for 25 minutes to remove any residual debris before analysis.

Extracts were analyzed within 24 hours by liquid chromatography coupled to a mass spectrometer (LC-MS). The LC–MS method was based on hydrophilic interaction chromatography (HILIC) coupled to the Orbitrap Exploris 240 mass spectrometer (Thermo Scientific).^50^ The LC separation was performed on a xBridge BEH Amide column (2.1 x 150 mm, 3.5 μm particle size, Waters, Milford, MA). Solvent A is 95%: 5% H_2_O: acetonitrile with 20 mM ammonium acetate and 20mM ammonium hydroxide, and solvent B is 90%: 10% acetonitrile: H_2_O with 20 mM ammonium acetate and 20mM ammonium hydroxide. The gradient was 0 min, 90% B; 2 min, 90% B; 3 min, 75% B; 5 min, 75% B; 6 min, 75% B; 7 min, 75% B; 8 min, 70% B; 9 min, 70% B; 10 min, 50% B; 12 min, 50% B; 13 min, 25% B; 14min, 25% B; 16 min, 0% B; 18 min, 0% B; 20 min, 0% B; 21 min, 90% B; 25 min, 90% B. The following parameters were maintained during the LC analysis: flow rate 150 mL/min, column temperature 25 °C, injection volume 5 µL and autosampler temperature was 5 °C. For the detection of metabolites, the mass spectrometer was operated in both negative and positive ion mode. The following parameters were maintained during the MS analysis: resolution of 180,000 at m/z 200, automatic gain control (AGC) target at 3 × 10^6^, maximum injection time of 30 ms and scan range of m/z 70-1000. Raw LC/MS data were converted to mzXML format using the command line “msconvert” utility.^51^ Data were analyzed using the EL-MAVEN software version 12. Student’s t-test was used to test for statistical significance, ** indicates p< 0.01; and *p< 0.05.

### LCMS Methods for Lipidomic Profiling

Tissue homogenization and extraction for lipids: Tissues were homogenized using a Retsch CryoMill. The homogenate was mixed with 1 mL of Extraction Buffer containing IPA/H2O/Ethyl Acetate (30:10:60, v/v/v) and Avanti Lipidomix Internal Standard (diluted 1:1000) (Avanti Polar Lipids, Inc. Alabaster, AL). Samples were vortexed and transferred to bead mill tubes for homogenization using a VWR Bead Mill at 6000 g for 30 seconds, repeated twice. The samples were then sonicated for 5 minutes and centrifuged at 15,000 g for 5 minutes at 4°C. The upper phase was transferred to a new tube and kept at 4°C. To re-extract the tissues, an additional 1 mL of Extraction Buffer (30:10:60, v/v/v) was added to the tube containing the tissue pellet. The samples were vortexed, homogenized, sonicated, and centrifuged as described earlier. The supernatants from both extractions were combined, and the organic phase was dried under liquid nitrogen gas.

Sample reconstitution for lipids: The dried samples were reconstituted in 300 µL of Solvent A (IPA/ACN/H2O, 45:35:20, v/v/v). After brief vortexing, the samples were sonicated for 7 minutes and centrifuged at 15,000 g for 10 minutes at 4°C. The supernatants were transferred to clean tubes and centrifuged again for 5 minutes at 15,000 g at 4°C to remove any remaining particulates. For LC-MS lipidomic analysis, 60 µL of the sample extracts were transferred to mass spectrometry vials.

LC-MS analysis for lipids: Sample analysis was performed within 36 hours after extraction using a Vanquish UHPLC system coupled with an Orbitrap Exploris 240™ mass spectrometer equipped with a H-ESI™ ion source (all Thermo Fisher Scientific). A Waters (Milford, MA) CSH C18 column (1.0 × 150 mm × 1.7 µm particle size) was used. Solvent A consisted of Acetonitrile (ACN): H_2_O (60:40; v/v) with 10 mM Ammonium formate and 0.1% formic acid, while solvent B contained Iso-propyl alcohol (IPA): ACN (95:5; v/v) with 10 mM Ammonium formate and 0.1% formic acid. The mobile phase flow rate was set at 0.11 mL/min, and the column temperature was maintained at 65 °C. The gradient for solvent B was as follows: 0 min 15% (B), 0–2 min 30% (B), 2–2.5 min 48% (B), 2.5–11 min 82% (B), 11–11.01 min 99% (B), 11.01–12.95 min 99% (B), 12.95–13 min 15% (B), and 13–15 min 15% (B). Ion source spray voltages were set at 4,000 V and 3,000 V in positive and negative mode, respectively. Full scan mass spectrometry was conducted with a scan range from 200 to 1000 m/z, and AcquireX mode was utilized with a stepped collision energy of 30% with a 5% spread for fragment ion MS/MS scan. Student’s t-test was used to test for statistical significance, ** indicates p< 0.01; and *p< 0.05.

### RNA Extraction

RT-qPCR was used to determine the change in mRNA levels in both human and mouse samples. In the human cohort, the two groups were a “young” cohort (18-55 years) and an “old” cohort (>60 years). In the mouse cohort, the “young” group consisted of mice aged 3 months, and the “old” group consisted of mice aged 2 years. Five samples were run for each experimental group for the human samples, and three samples were run for each experimental group for the mouse samples. For both experimental groups, comprising both human and mouse samples, at least three samples were necessary for a statistical test to determine significant differences.

Murine liver tissue was collected via microdissection, while the human tissue was collected by macrodissection. Collected samples were immediately placed in liquid nitrogen to freeze and were stored in -80°C for 3 months prior to initiation of reverse transcription and qPCR. To isolate RNA, we first used the TRIzol reagent (Invitrogen) protocol on its user guide, with further purification of isolated RNA being performed with the RNeasy Micro kit and protocol (Qiagen Inc), including DNase treatment of RNA samples with kit-included DNase I. RNA concentration was determined by measuring absorbance at 260 nm and 280 nm using a NanoDrop 1000 spectrophotometer (NanoDrop products, Wilmington, DE, USA). The determination of significant DNA contamination was assessed by the A280/A260 ratio, with a ratio of 1.8-2.0 being necessary for RNA samples to be considered sufficiently pure for downstream applications.

### RT-qPCR

Reverse transcription was performed with the High-Capacity cDNA Reverse Transcription Kit (Applied Biosciences, Carlsbad, CA). A total reaction volume of 20 μL, containing 1 μg of RNA, was used. The kit-included MultiScribe™ Reverse Transcriptase (2.5 U/μL) was used as the reverse transcriptase. We used the thermal cycle recommended by the kit’s protocol: 25°C for 10 minutes, 37°C for 120 minutes, 85°C for 5 minutes, and then held at 4°C.

Quantitative PCR (qPCR) was then performed using SYBR Green (Life Technologies, Carlsbad, CA).^52^ For qPCR, a 20 μL reaction volume containing 50 ng of cDNA was loaded into each well of a 384-well plate for nuclear gene expression measurement, with the reaction carried out on an ABI Prism 7900HT system (Applied Biosystems). For qPCR for mtDNA, five nanograms of DNA were used for the quantification of mitochondrial (Cox1) and nuclear (Rpl13a) DNA markers with Rpl13a. The reaction was performed using primers (Table 1) at a concentration of 500 nM and with dNTP provided by 2X SYBR^®^ Green PCR Master Mix diluted to 1X (Life Technologies). The DNA polymerase used was kit-included AmpliTaq Gold^®^ DNA polymerase, diluted by the recommended factor from 2X to 1X. The SYBR^®^ Green PCR Master Mix also included the buffer necessary for the protocol. Thermal cycling parameters were adapted from previous qPCR protocol^45^ with the following cycle: 1 cycle at 95°C for 10 min; 40 cycles of 95°C for 15 s; 59°C for 15 s, 72°C for 30 s, and 78°C for 10 s; 1 cycle of 95°C for 15 s; 1 cycle of 60°C for 15 s; and one cycle of 95°C for 15 s. GAPDH normalization was used to present the data as fold changes. qPCR primers used were from previously published sequences,^45^ as detailed in **Table 1**.

Specificity for fragments was determined by matching amplicon sizes, as determined by gels with positive control samples. For no template controls (NTCs), detection of DNA amplification remained negative after 40 cycles. Quantification of Ct for runs was determined by fitting to a 6-point standard curve of log DNA dilution vs. Ct (y-intercept: 38.664; slope: -3.57; r^2^=.96; PCR efficiency: 90.5%). Cq variation at the lower limits was minimal, confirming the assay’s accuracy. The limit of detection was determined by the melting curve.

qPCR analysis was performed using the RQ Manager software on the ABI Prism 7900HT system (Applied Biosystems), which was utilized for qPCR. The Automatic C_T_ setting on RQ Manager was used to determine the C_T_ for samples, with outliers determined by using the C_T_ vs. Well Position View plot on the RQ Manager software. GAPDH was used as the sole reference gene due to its widespread use as a reference gene across multiple protocols^53^. A comparison of results across treatments was performed by normalizing target gene expression to GAPDH. Intra-assay variation for samples of the same treatment, run on the same plate, was measured to ensure sufficient repeatability in the experiment. For each reverse transcription and qPCR experiment, there were 3 technical replicates. To determine significance, Mann–Whitney tests were performed with an α of 0.05, and the results were visualized in GraphPad Prism 10.2.3 (La Jolla, CA, USA).

### Western Blotting

Western blotting was performed as previously described.^54^ Briefly, following homogenization and lysis in RIPA lysis buffer (1% NP40, 150 mM NaCl, 25 mM Tris base, 0.5% sodium deoxycholate, 0.1% SDS, 1% phosphatase inhibitor cocktails #2 (Sigma P5726-1ML) and #3 (Sigma P0044-1ML), and one complete protease inhibitor tablet (Sigma 04693159001)), protein concentration in 3-month and 2-year tissue lysates were quantified using a BCA Assay (Thermo Scientific VLBL00GD2). Equal amounts of proteins were run on 4%–20% Tris-glycine gels (Invitrogen WXP42012BOX). Protein was then transferred to a nitrocellulose membrane (Li-Cor 926-31092) and incubated overnight at 4°C with gentle agitation in the presence of primary antibodies: Mic60/mitofilin (Abcam ab110329), SAM50 (Proteintech 20824-1-AP), or tubulin (Novus NB100-690). Secondary antibodies [1:10,000; donkey anti-mouse IgG (H + L) (Invitrogen A32789) and donkey anti-rabbit IgG (H + L) (Invitrogen A32802)] were incubated with the membrane at room temperature for 1 h. Using the Li-Cor Odyssey CLx infrared imaging system, blots were imaged. We performed the student’s t-test for statistical significance, with ** indicating p < 0.01 and *indicating p < 0.05.

### Confocal mCherry-Mito-7 Labeling

To label the mitochondria of cardiac fibroblasts, the mCherry-Mito-7 plasmid was transfected into the cells using a transfection reagent according to the manufacturer’s instructions^55^ and as previously described.^80^ Briefly, following plasmid and transfection reagent dilution in Opti-MEM medium and incubation at room temperature for 20 minutes, the dilution was added to the culture medium of the cells, which were incubated for 24–48 hours to allow expression of the mCherry-Mito-7 protein. Localization in fibroblasts was visualized using a Leica SP8 Confocal Microscope. Cells were washed with PBS, fixed with 4% paraformaldehyde for 10 minutes, and mounted with DAPI-containing mounting medium. Fluorescent signals were observed using appropriate filters and recorded with a digital camera.^31,36–40,56–59^

### Knockdown of *MIC60* and *CHCHD6* in HepG2 cells

The transfection of *MIC60* and *CHCHD6* siRNAs into HepG2 cells was carried out using Lipofectamine RNAiMax (Invitrogen) in accordance with the manufacturer’s instructions and as previously described.^60^ Authentication of cell line determined by confirming proper morphology by microscopy. Following a 48-hour incubation period, the cells were utilized for mitochondrial calcium (_m_Ca^2+)^ and ROS measurements.

### Measurement of mitochondrial calcium uptake and retention capacity in HepG2 cells

The _m_Ca^2+^ uptake and retention capacity in HepG2 cells were assessed using a ratiometric Ca^2+^ sensor Fura-FF, as detailed earlier,^61^ with slight modifications. In brief, cells (2.5x10^6^) were washed with Ca^2+^/Mg^2+^-free DPBS (GIBCO), permeabilized in intracellular medium (ICM: 120 mM KCl, 10 mM NaCl, 1 mM KH_2_PO_4_, 20 mM HEPES-Tris, pH 7.2), and supplemented with thapsigargin and digitoxin. Fura-FF (1 μM) was added at the 0s time point, and fluorescence emissions at 340- and 380 nm excitation/ and emission at 510 nm were monitored using a multi-wavelength excitation dual-wavelength emission fluorimeter (Delta RAM, PTI). To assess the _m_Ca^2+^ uptake, a bolus of 5 μM Ca^2+^ and the mitochondrial uncoupler FCCP (10 μM) were added at specified time points with continuous stirring at 37°C. To assess mCa^2+^ retention capacity, following baseline recordings, a series of Ca^2+^ boluses (5 µM) were introduced at specified time points. Upon reaching a steady state, 10 μM FCCP was added to collapse the Δψm and release matrix free-Ca^2+^. The number of Ca^2+^ boluses taken up by cells was counted to calculate mitochondrial CRC. For all statistical tests, a one-way ANOVA statistical test was performed with Dunnett’s multiple comparisons test. N = 5 -10 for all calcium experiments, each indicated by dots, as run in triplicate. Significance values indicate **P ≤ 0.01 and ****P ≤ 0.0001.

### Evaluation of ROS levels

Approximately 0.2 million HepG2 cells were plated in 35 mm dishes. The next day, MIC60 and CHCHD6 siRNAs were transfected using Lipofectamine RNAiMax (Invitrogen) according to the manufacturer’s instructions. After incubation for 30 hrs., cells were co-stained for 30 minutes at 37°C with two different dyes for ROS detection: MitoBright ROS Deep Red (10 µM, Dojindo Laboratories) for mitochondrial superoxide, and DCFDA (10 µM, Invitrogen) for intracellular total ROS. Following incubation with staining dyes, cells were washed three times with 1X HBSS, and ROS analysis was performed using a confocal microscope (FV4000, Olympus Life Science).

For mitochondrial H_2_O_2_ assessment, cells were stained with MitoPY1 (5 µM, Bio-Techne) for 45 min at 37°C. Cells were then washed with 1x HBSS and imaged using a confocal microscope (FV4000, Olympus Life Science). ImageJ was used for quantifying fluorescence intensities. For all statistical tests, one-way ANOVA statistical test was performed with Dunnett’s multiple comparisons test. N = 9 -13 for all oxidative stress experiments, each indicated by dots, as run in triplicates. Significance values indicate **P ≤ 0.01 and ****P ≤ 0.0001.

### Data Analysis

GraphPad Prism 10.2.3 (La Jolla, CA, USA) was used for all statistical analyses. All experiments involving SBF-SEM and TEM data had at least three independent experiments. Statistics were not handled by those conducting the experiments. For all experiments, at least three samples were used for each experimental group to enable statistical testing. The black bars represent the standard error of the mean. For all analyses, one-way ANOVA was performed with tests against each independent group and significance was assessed using Fisher’s protected least significant difference (LSD) test. *, **, ***, **** were set to show significant difference, denoting *p* < 0.05, *p* < 0.01, *p* < 0.001, and *p* < 0.0001, respectively.

## Results

### Genetically Regulated MICOS Gene Expression Associated with Liver Disease Across Ancestries

GREX models derived from GTEx v8 were applied to the BioVU cohort, enabling systematic association testing between predicted expression levels of MICOS genes responsible for cristae stabilization, formation, remodeling and cell survival, and liver disease phenotypes across various tissues (Figure 1A). To examine the relationship between MICOS complex genes *CHCHD3 (MIC19), CHCHD6 (MIC25), and OPA1* and liver disease, we leveraged Vanderbilt University Medical Center’s EHR-linked biobank, BioVU (**Table 2**). We used logistic regression analysis to test whether the GREX of *CHCHD3, CHCHD6, and OPA1* was associated with liver disease phenotypes extracted from the EHR (Figure 1B-E). Although no associations met a strict global Bonferroni-corrected p-value threshold, we found significant associations that met the less stringent within-tissue Bonferroni-corrected p-value (see Methods). In individuals of European genetic ancestry (EUR), we identified significant associations between decreased GREX of *CHCHD3* in three tissues with liver transplant and liver cirrhosis without mention of alcohol (p < 0.0056, Figure 1B). In individuals of African genetic ancestry (AFR), we identified significant associations between decreased GREX of *OPA1* in five tissues and chronic liver disease and cirrhosis (p <0.0056, Figure 1E).

**Figure 1.**
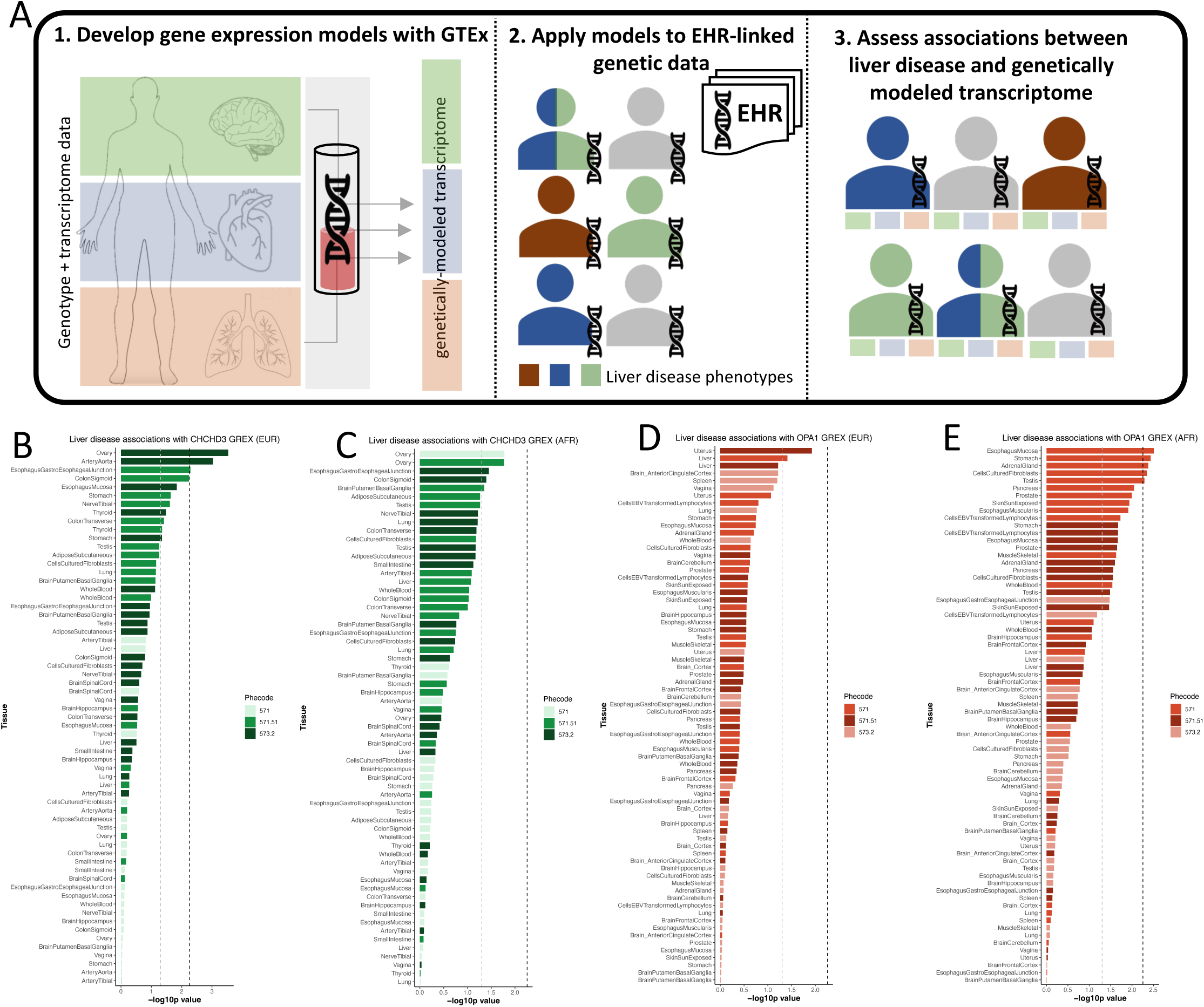
**Assessing the relationship between *CHCHD3, CHCHD6*, and *OPA1*** genetically modeled gene expression and liver diseases in a clinical biobank. (A) Genetically regulated gene expression (GREX) was modeled from GTEx version 8 data, which includes genotype data linked to measured gene expression across 49 tissues. GREX was calculated within the BioVU population for the MICOS complex genes CHCHD3, CHCHD6, and OPA1. Liver disease phenotypes were extracted from the de-identified medical records of BioVU participants. Associations between the MICOS GREX and liver phenotypes were performed using logistic regression models. (B-C). Association results for CHCHD3 GREX in (B) 65,363 individuals of European (EUR) genetic ancestry and (C) 12,313 individuals of African (AFR) genetic ancestry with three liver disease phenotypes. (D-E) Association results for OPA1 GREX in (D) 65,363 individuals of European (EUR) genetic ancestry and (E) 12,313 individuals of African (AFR) genetic ancestry with three liver disease phenotypes (phecode 571: Chronic liver disease and cirrhosis, phecode 571.51: Cirrhosis of liver without mention of alcohol, phecode 573.2: Liver replaced by transplant). Grey line annotates nominal significance (p=0.05) and black line annotates the within-tissue Bonferroni-corrected p-value (p=0.0056). GREX tissue model results are ranked by -log10p value on the y axis and -log10p values are represented on the x axis. A comprehensive supplementary table with all association results (effect sizes, standard errors, p-values) for all gene-tissue-phenotype combinations are found in the supplementary information.

### Aging is Associated with Coordinated Suppression of MICOS Genes and Activation of ER Stress Pathways

To further investigate whether the MICOS genes that regulate mitochondrial bioenergetics, ER stress, and apoptosis differ with aging, we performed qPCR in both human (Figure 2A-I) and mouse (Figure 2J-M) and compared the results between young and old. In the human cohort (n = 5/group), the expression of MICOS proteins *Mic19, Mic25, And Mic60* (Figure 2A, B, C) was reduced in the older compared to the younger samples. Genes associated with regulating mitochondrial dynamics and tethering to the ER, such as *BIP* (Figure 2D), *PERK* (Figure 2E), *XBP1* (Figure 2F), *CHOP* (Figure 2G), *IRE1* (Figure 2H), and *ATF4* (Figure 2I), had increased expression. In the murine model (n = 3/group), we observed reduced expression of MICOS proteins including, *Opa1, Mitofilin, Chchd3, and Chchd6* (the murine ortholog of *MIC60*), in old mice compared with young mice (Figure 2J-M).

**Figure 2:**
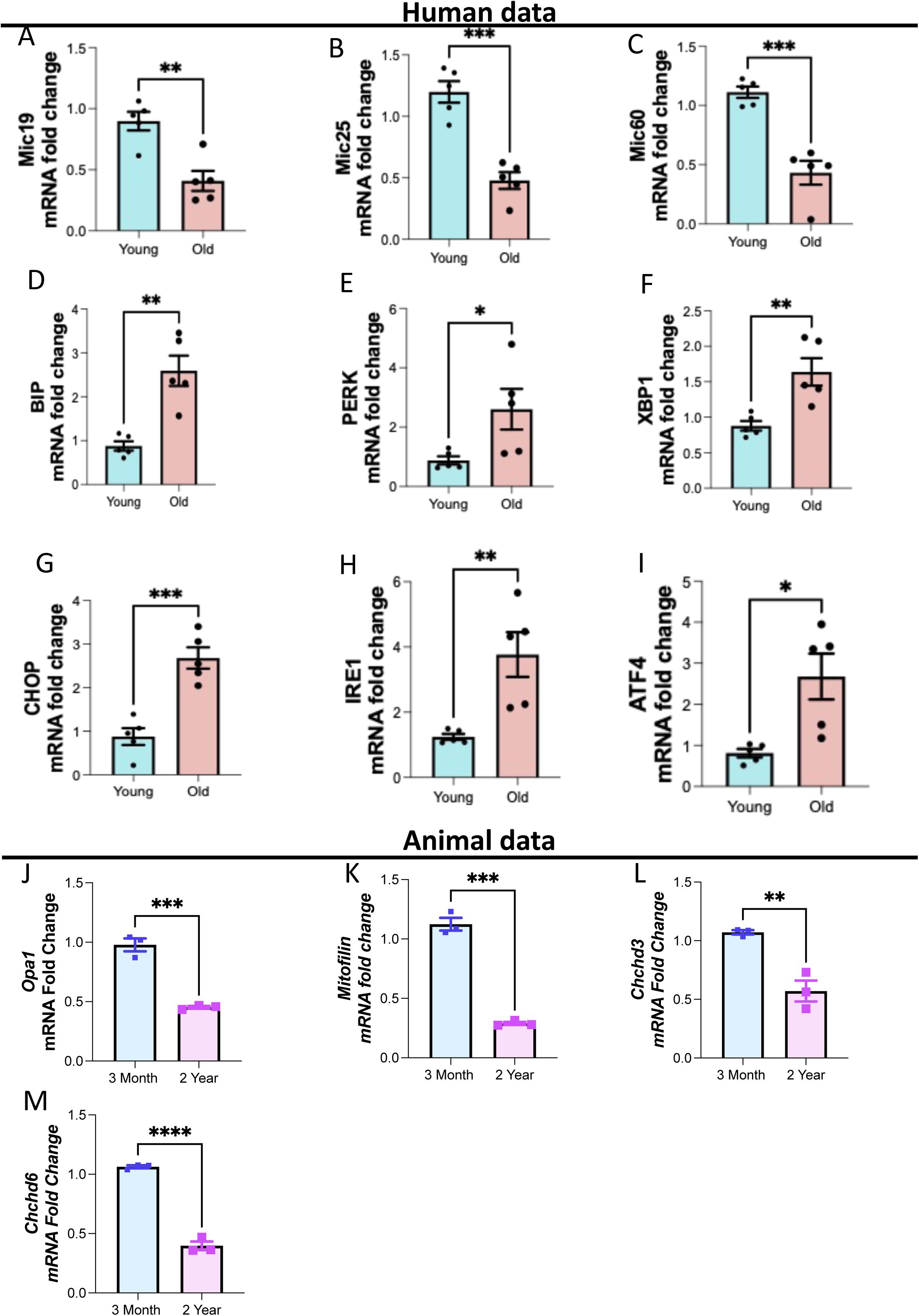
MICOS mRNA fold changes in young and old subjects. Human mRNA fold change in (A) Mic19, (B) Mic25, (C) Mic60, (D) BIP, (E) PERK, (F) XBP1, (G) CHOP, (H) IRE1, and (I) ATF4 expressions between young and old human liver tissue samples (n = 5/group). Murine model mRNA fold change in (J) Opa1, (K) Mitofilin, (L) Chchd3, and (M) Chchd6 expression in mouse liver tissue between young (3 months of age) and old (2 years of age) cohorts (n = 3/group). Mann–Whitney tests were used for statistical analysis. Statistical significance is denoted as ns (not significant), *p < 0.05, **p < 0.01 and ***p < 0.001.

### Loss of MICOS Components Disrupts Calcium Homeostasis and Promotes Cellular Stress in HepG2 Cells

While the reduction in expression of *Mic19, Mic25*, and *Mic60* highlights compromised mitochondrial bioenergetics, and the increased expression of Bip, Xbp1, Ire1, Atf4 and Chop may suggest proteostatic and metabolic stress. We sought to demonstrate and investigate the role of the MICOS complex in age-related changes by modeling MICOS loss *in vitro*. To investigate the role of *MIC60* and *CHCHD6* in mitochondrial calcium (_m_Ca^2+^) homeostasis, we monitored _m_Ca^2+^ uptake and _m_Ca^2+^ retention capacity (CRC) in human-derived HepG2 cells with *MIC60* and *CHCHD6* knockdown (Figure 3). Raw mitochondrial calcium uptake traces showed reduced calcium uptake dynamics in permeabilized HepG2 cells following *MIC60* or *CHCHD6* knockdown compared with scramble-siRNA controls (Figure 3A and B). Furthermore, to determine if the altered MICOS and cristae structure contributes to impaired CRC and is involved in mitochondrial permeability transition pore opening, we measured the CRC in *MIC60* and *CHCHD6-knockdown* cells. Knockdown of Mic60 as well as Chchd6 sensitized cells to mPTP opening, as indicated by a sudden Ca^2+^ release at a lower cumulative Ca^2+^ load compared to scramble-siRNA controls (Figure 3C). We also found that knocking of *MIC60* and *CHCHD6* in HepG2 cells significantly reduced _m_Ca^2+^ retention capacity (Figure 3D). Quantitative analysis confirmed a significant reduction in CRC in both knockdown conditions relative to scramble-siRNA–treated cells (Figure 3D). Immunoblot analysis verified efficient siRNA-mediated knockdown of Chchd6 (Figure 3E) and Mic60 (Figure 3F) in HepG2 cells. These data suggest that altered MICOS complex and cristae disorganization increase the susceptibility of HepG2 cells to _m_Ca^2+^ dysregulation and Ca^2+^-induced cell death.

**Figure 3:**
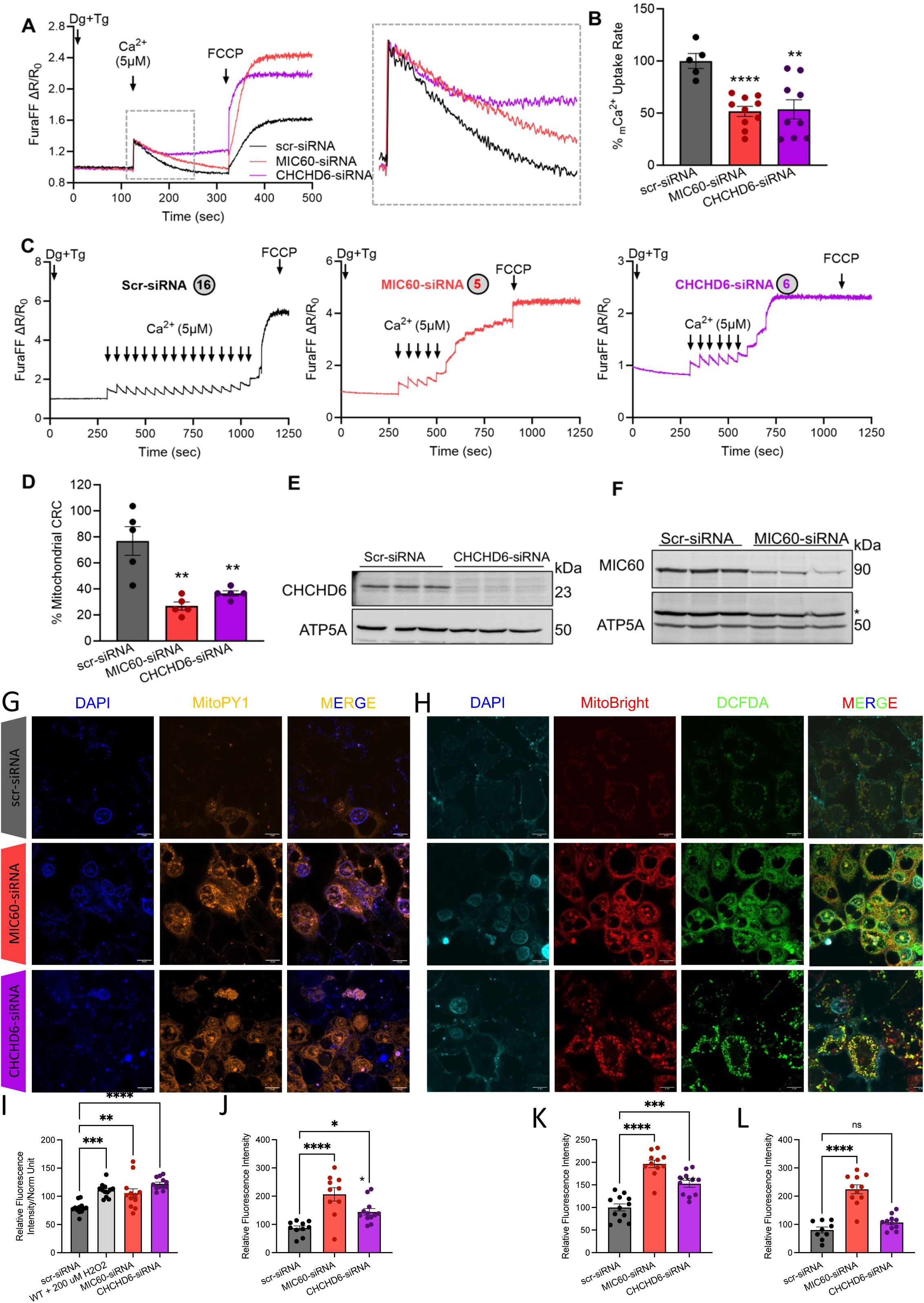
Loss of *Mic6*0 and *Chchd6* in HepG2 cells results in reduced mCa2+ uptake and calcium retention capacity and oxidative stress. (A) Raw traces showing mitochondrial calcium uptake in permeabilized *MIC60* and *CHCHD6* knockdown HepG2 cells along with scr-siRNA transected controls. (B) Percentage change in mCa2+ uptake rate quantified from raw traces. (C) Recordings of mitochondrial calcium retention capacity in scr-siRNA, *MIC60* siRNA, and *CHCHD6* siRNA HepG2 cells. The circles indicate the number of calcium boluses taken up by specific cells. (D) Percentage change in mitochondrial calcium retention capacity quantified from recordings of mitochondrial calcium retention capacity (CRC). (E) Immunoblot confirming siRNA-mediated knockdown of *CHCHD6* in HepG2 cells. (F) Immunoblot confirming siRNA-mediated knockdown of *MIC60* in HepG2 cells. (G) 4′,6-diamidino-2-phenylindole (DAPI) staining, MitoPY1 (5 µM, 45 min at 37°C magnification of 60x), and merge channels in scramble-siRNA (control), MIC60-siRNA, and CHCHD6-siRNA transfected permeabilized HepG2 cells. (H) 4′,6-diamidino-2-phenylindole (DAPI) staining, MitoBright Deep Red (10 µM, 30 min at 37°C), DCFDA (10 µM, 30 min at 37°C, magnification of 60x), and merge channels in scramble-siRNA (control), MIC60-siRNA, and CHCHD6-siRNA transfected permeabilized HEK293 cells. (I) Plate reader-based reactive oxygen species (ROS) quantification. (J) Microscopy-based ROS quantification of MitoPY1 orange, (K) MitoSox Deep Red, and (L) DCFDA. For all statistical tests, a one-way ANOVA statistical test was performed with Dunnett’s multiple comparisons test. N=5-10 for all calcium experiments, each indicated by dots, as run in triplicate. N=9-13 for all oxidative stress experiments, each indicated by dots, as run in triplicate. Significance values indicate **P ≤ 0.01 and ****P ≤ 0.0001.

Since Ca^2+^ directly affects oxidative stress signaling and the generation of reactive oxygen species (ROS),^81^ we wanted to determine if MICOS influences ROS production. We evaluated total ROS, mitochondrial superoxide, and H_2_O_2_ production in *MIC60* and *CHCHD6* knockdown cells (Figures 3G and H). Mitochondrial H_2_O_2_ content, as measured by Mitochondria Peroxy Yellow 1 (Figure 3G), increased following *MIC60* and *CHCHD6* knockdown when quantified by both plate-reader-based (Figure 3I) and microscopy-based (Figure 3J) ROS quantification. Silencing *MIC60* and *CHCHD6* in HepG2 cells significantly increased mitochondrial superoxide production and more general intracellular ROS, detected by MitoBright Deep Red (Figure 3K) and DCFDA (Figure 3L), respectively. These findings indicate that suppression of *MIC60* and *CHCHD6* disrupts mitochondrial ROS homeostasis, demonstrating that the MICOS complex is associated with oxidative stress.

### Aging is Associated with Broad Alterations in Liver Metabolism, Including Changes in Energy Utilization and Storage

Due to observed changes in liver gene expression and function during aging, we proceeded to analyze alterations in small biomolecules that regulate energy metabolism, cellular homeostasis, and storage. Our investigation involved global metabolic profiling of young (n = 4) and aged (n = 4) liver tissues, revealing significant changes in aged mouse livers (Figure 4A and B). Principal component analysis showed a clear separation between the metabolic profiles of young and aged livers (Figure 4C). We noted an accumulation of metabolites related to Vitamin A metabolism, specifically retinoic acid and retinol (Figures 4D and E). Our metabolomics analysis also uncovered an age-related decrease in tricarboxylic acid (TCA) cycle intermediates: succinate and malate, which could be consistent with increasing cataplerosis associated with aging for biosynthetic use or impaired activity (Figures 4F and G). Furthermore, we observed decreases in nucleotide monophosphates involved in purine and pyrimidine metabolism, including GMP, CMP, UMP, and AMP (Figures 4H-K). The synthesis of these nucleotides, which involves mitochondrial processes, is influenced by age-related mitochondrial dysfunction, thereby affecting the overall nucleotide biosynthesis pathway, which supports our observations^66^. Our results additionally confirmed existing literature on dysregulated nicotinamide adenine dinucleotide (NAD^+^) metabolism in the aging liver (Figure 4M).^63^ Significant depletions in tissue ADP, NAD^+^, NADP, and NMN pools were detected (Figure 4L-O). In summary, these findings collectively support the presence of altered energy metabolism and cellular homeostasis in the metabolically active liver, resulting from mitochondrial dysfunction and impairment.

**Figure 4.**
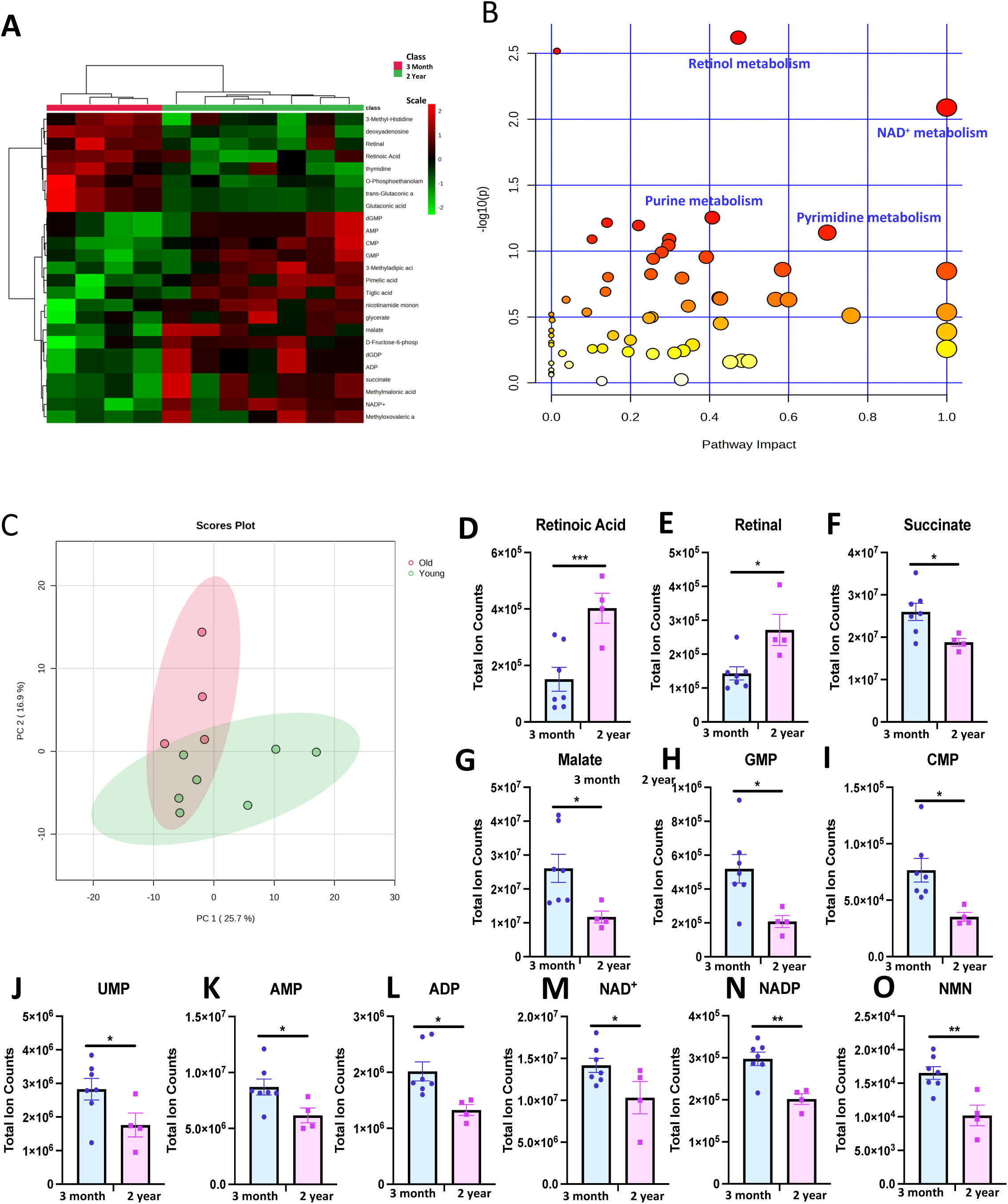
Global metabolomic profiling reveals changes in lipid composition, mitochondrial DNA, and metabolic pathways. (A) Metabolomic heatmap of the top 25 most enriched metabolic pathways in 3-month and 2-year mouse liver tissue. For each tissue and metabolite in the heatmaps, the 2-year samples were normalized to the median of the 3-month samples and then log2 transformed. (note: p-values were adjusted to correct for multiple comparisons using an FDR procedure) and log fold changes greater than 1 or less than −1. 3-month, n= 4; 2-year, n= 4. (B) Pathway impact graph of metabolic pathways in 3-month and 2-year mouse liver tissue that are affected by aging. A higher impact factor and a larger circular area are associated with greater fold enrichment of metabolic pathways. More red coloring of circles is associated with higher statistical significance. (C) PCA chart of metabolite concentration between 3-month and 2-year-old samples. (D-O) Concentration of metabolites (D) retinoic acid, (E) retinal, (F) succinate, (G) malate, (H) GMP, (I) CMP, (J) UMP, (K) AMP, (L) ADP, (M) NAD+, (N) NADP, and (O) NMN in mouse liver in 3-month (blue) and 2-year-old (pink) samples. For all panels, error bars indicate SEM, Mann–Whitney tests were used for statistical analysis, and significance values indicate *P ≤ 0.05, **P ≤ 0.01, ***P ≤ 0.001, ****P ≤ 0.0001, and ns indicates non-significant. For all panels, error bars indicate SEM, ** indicates p< 0.01; and *p< 0.05, calculated with Student’s t-test.

### Aging Causes Alterations in Liver Lipid Accumulation in humans and mice

Based on the evidence of an altered metabolomic profile in aged livers, we also hypothesized that lipid metabolism is altered in aged livers. Damage to the liver’s tissue and alterations in hepatic metabolism during aging might result from abnormal lipid levels in the liver.^18^ First, we utilized magnetic resonance imaging (MRI) to determine how lipid content changes during aging in human samples. By enrolling female and male participants (n = 10 per group) (Figures 5A–D’), we created a “young” cohort consisting of individuals from 18 to 55 years old and an “old” cohort of individuals older than 60 years old. For both sexes, the percentage of lipid in the liver was higher in the old compared to the young (Figures 5E and F). Specifically, when combining the male and female cohorts (Figure 5G), the mean fat percentage was higher in the older group (5.93 ± 6.11% SD) compared to the younger group (1.41 ± 1.44% SD). Additionally, the 75% percentile of fat percentage was higher, at 10.5% in the aged cohort, as compared to 2.10% in the young cohort. Notably, Grade 1 of fat fraction classification, representing mild hepatic steatosis, has a fat percentile cutoff of 6.5% or higher.^42^

**Figure 5.**
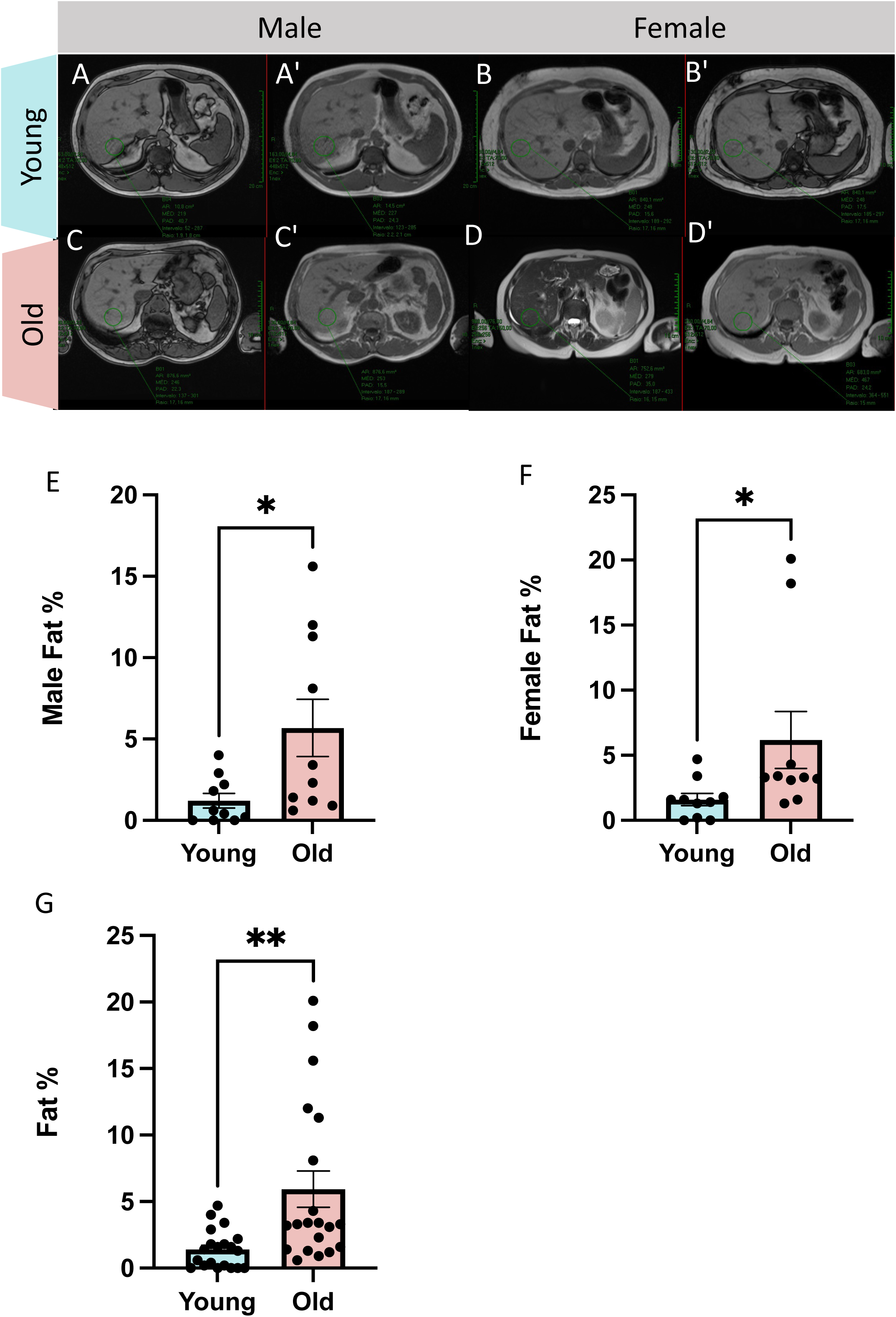
Comparative analyses of liver lipid content in young and old humans (A-D’) Cross-sectional imaging of in-phase (A-D) and out-of-phase (A’-D’) liver anatomy data from (A,A’) males under 55 years old (aged 14–53 years old; n = 10), (B,B’) females under 55 years old (aged 17–41 years old; n = 10), (C,C’) males over 60 years old (aged 60–85 years old; n = 10), and (D,D’) females over 60 years old (aged 64–96 years old; n = 10). (E-F) Liver lipid percentage in young and old cohort in (E) males and (F) females and (G) Body Fat%.

To further assess the intricate changes in the lipid profile, we used murine livers. We used Oil Red O, a neutral lipid stain, to visualize lipids in 3-month and 2-year murine liver tissue (Figures 6A-D). We saw an increased amount of Oil Red O in the 2-year cohort (n=10 per group), signifying an increase in lipid amount across the aging process in the liver, with a more than 6-fold increase in the number of lipid droplets in 2-year samples (Figure 6E; 3-months: 12.2 ± 6.55 SD; 2-years: 77.5 ± 20.4 SD). An uptick of lipid droplets in hepatocytes is indicative of MASLD.^65^ To further explore markers of potential disease progression and dystrophy outside of lipid accumulation, we also looked at liver weight, which was decreased in aged samples when normalized to body weight (Figure 6F; 3-months: 4.56 ± 0.329% SD; 2-years: 3.93 ± 0.376% SD). Since mtDNA content reduction is another hallmark of MASLD, we also measured mtDNA content and observed an age-related decline (Figure 6G; 3 months: 1.00 (normalized mean); 2 years: 0.805 ± 0.119 SD). Additionally, abnormal liver function and disease states increase bile acid concentrations, which the liver normally metabolizes.^67^ We looked at the concentration of bile acid, but no significant difference was noted (Figure 6H; 3 months: 0.871 ± 0.138 µMol/µg SD; 2 years: 0.89 ± 0.15 µMol/µg SD). Finally, central to the pathology and a key marker of MASLD is high triglycerides.^68,69^ We observed that aging murine samples had significantly elevated triglycerides in the liver (Figure 6I; 3 months: 0.30 ± 0.10 mmol/L SD; 2 years: 0.748 ± 0.15 mmol/L SD), which was more drastic than the general increase in serum triglycerides (Figure 6J; 3 months: 0.211 ± 0.100 mmol/L SD; 2 years: 0.379 ± 0.118 mmol/L SD). TEM analysis revealed increased lipid droplet accumulation in livers from 2-year-old mice compared with 3-month-old controls (Figure 6K and L). Aged samples had significantly increased both the number of lipid droplets per unit area and the total lipid droplet area within TEM images (Figure 6M and N), indicating enhanced hepatic lipid deposition with age. These results support the observation of an age-related progression toward liver disease through the accumulation of hallmarks associated with MASLD.

**Figure 6.**
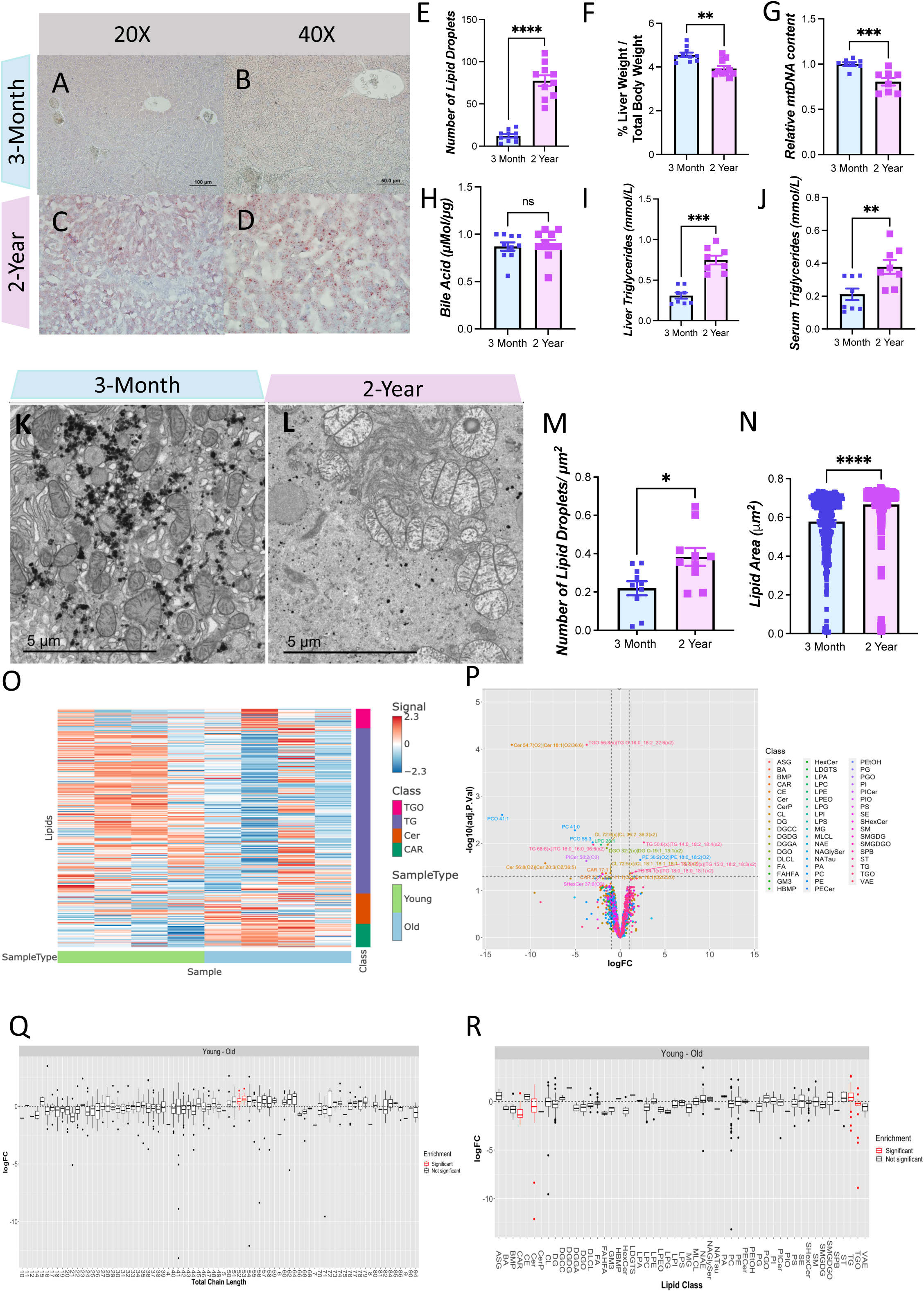
Global lipidomic profiling reveals variations in lipid classes and chain lengths with age in liver tissues. (A-D) Oil Red O Staining of liver tissue from 3-month (A-B) and 2-year (C-D) old mice at (A, C) 20x and (B, D) 40x magnification. (E) Quantification of Oil Red O staining per area (n=10 per group). (F) Quantification of liver weight relative to total body weight as a percent (n=10), (G) relative real-time quantitative polymerase chain reaction mtDNA content (n=8), (H) bile acids concentrations (n=10), (I) liver triglycerides concentration (n=8), (J) and serum triglycerides concentration (n=8). (K-N) TEM image of mouse liver tissue from (K) 3-month and (L) 2-year-old mice. (M) Number of lipid droplets per μm^2^ of area in TEM image. (N) Total area of TEM image consisting of lipid droplets in μm^2^. (O) Lipid class enrichment heatmap. (P) Volcano plot labeling significant hits in lipid expression between 3-month and 2-year-old livers, which have adjusted p-value <0.05 and fold change (+ or -) greater than 1. (Q) Lipid class (R) and lipid chain length enrichment based on comparison between 3-month and 2-year-old livers. For each tissue and metabolite in the heatmaps, the 2-year-old samples were normalized to the median of the 3-month samples and then log2 transformed. Significantly different lipid classes represented in the figures are those with adjusted p-values < 0.05 (note: p-values were adjusted to correct for multiple comparisons using an FDR procedure) and log fold changes greater than one or less than −1. 3-month, n= 4; 2-year-old, n= 4. For all panels, error bars indicate SEM, ** indicates p< 0.01; and *p< 0.05, calculated with Student’s t-test.

Lipidomic profiling of both young and aged liver tissues unveiled age-related changes in lipid classes and chain lengths (Figure 6O-R). Lipid class enrichment heatmap and volcano plot analyses highlighted distinct age-dependent shifts in lipid abundance (Figures 6O and P). In the aging liver, we observed significant alterations in triglyceride oligomers (TGO), triglycerides (TG), ceramides (Cer), and acylcarnitines (CAR) relative to other lipid classes (Figure 6Q). Aging also significantly altered lipid chain-length distributions in the liver (Figure 6R), suggesting changes in membrane composition that may influence membrane integrity, fluidity, and function. Together, these findings align with the altered metabolomic and lipogenic remodeling associated with age-related changes in mitochondrial structure and function.

### Ultrastructural Changes in Murine Liver Reveal Aging is Associated with Lower Mitochondrial Volume and Complexity

Next, we sought to determine whether these age-related changes in liver mass, disease markers, and metabolomic and lipidomic profiles are correlated with atypical mitochondrial structures in the liver. For these studies, we utilized aged C57BL/6J mice at 2 age points, 3 months, and 2 years, which are generally understood to be good representations of “young” and “old” aged points akin to aging displayed in humans.^74^ To begin with, we used TEM to measure changes in mitochondrial morphology in aged mice (Figures 7A-D). TEM images revealed an increase in lipid droplets in the livers of aged individuals of both sexes, which appeared less circular. TEM analysis revealed that the mitochondrial count increased while the average mitochondrial area decreased with aging (Figures 7E-H). Analysis by two-way ANOVA (Age versus Sex) revealed a significant main effect of age on mitochondrial count (Figure 7I) and area (Figure 7J), but no significant interaction with sex, indicating similar age-related trends in both males and females. In aged male and female murine samples, we found a significant decrease in cristae score (a semi-quantitative measure of cristae integrity^14^) in both sexes (Figures 7K and L). A two-way ANOVA confirmed a main effect of age (Figure 7M). Together, these results highlight that cristae integrity is lower in aged liver tissue samples, indicating a loss of ATP production efficiency. Given these findings and the resource-intensive nature of 3D imaging, we focused the subsequent SBF-SEM analysis on male mice (Figure 7N).

**Figure 7:**
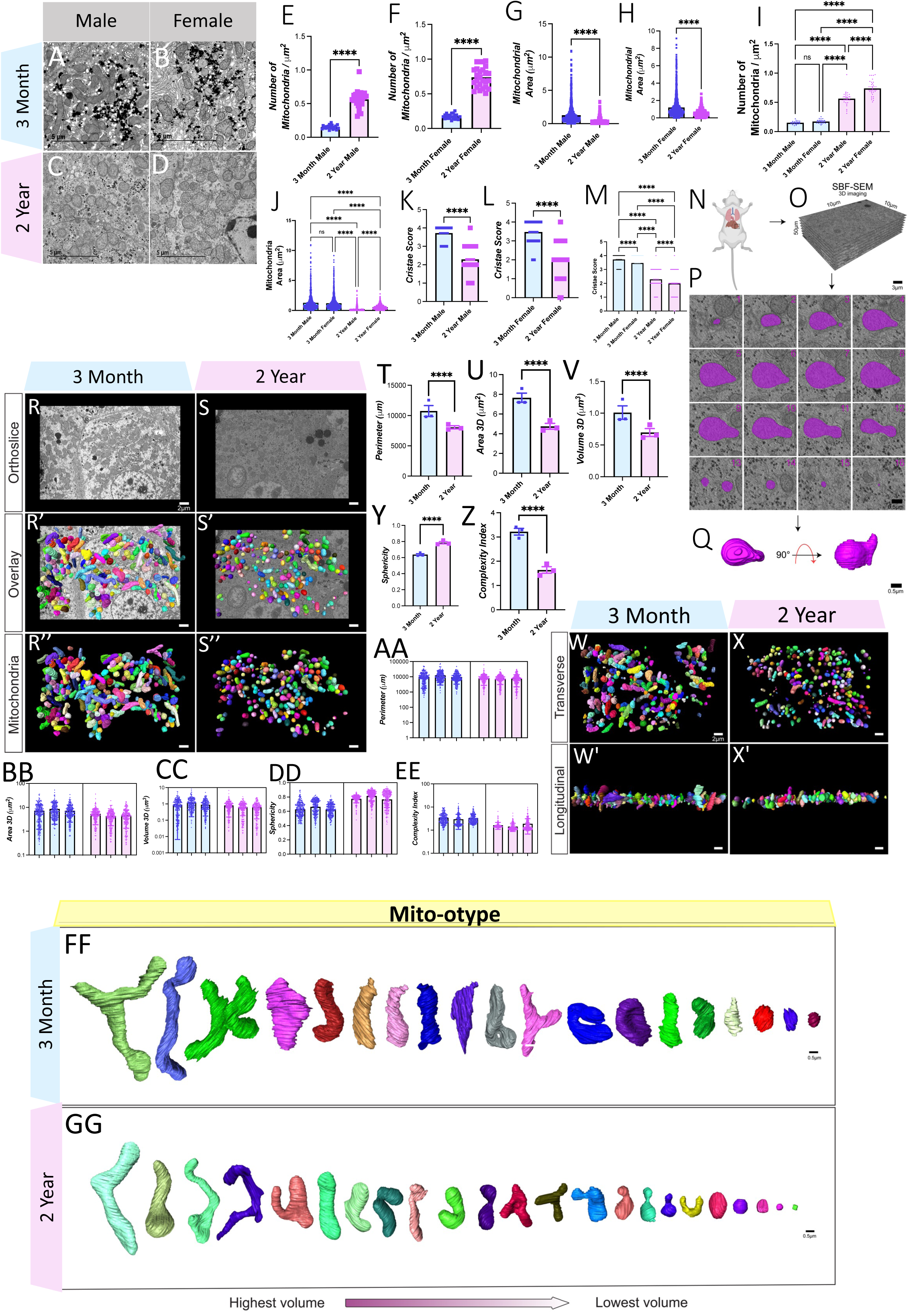
TEM and SBF-SEM Shows Changes in Mitochondria Murine Liver Across Aging. (A-D) Representative transmission electron micrographs in (A, C) males and (B, D) females from (A-B) 3-month and (C-D) 2-year murine liver tissue. (E) Mitochondrial quantifications of male mitochondria number per μm^2^ area in TEM image (n=21 images across 5 mice, 3-months; n=24 images across 5 mice, 2-years) (G) mitochondrial area quantified in 3 mice, over 3 ROI (n=1309 mitochondria, 3-months; n=1333, 2-years) and (K) cristae score (n=555, 3-months; n=555, 2-years) in male murine liver TEM images. (F) Mitochondrial quantifications of the number of female mitochondria per μm^2^ unit area in the TEM image. (n=21, 3-months; n=25, 2-years) (H) mitochondrial area (n=1253, 3-months; n=1018, 2-years) (I) Combined Mitochondrial quantifications of male and female mitochondria number per μm^2^ unit area in TEM image across aging. (n=21 each sex, 3-months; n=25 each sex, 2-years) (J) Combined mitochondrial area for males and females across aging (n=1253 female, n=1309 male, 3-months; n=1018 female, n=1333 males, 2-years) and (L) cristae score (n=684, 3-months; n=684, 2-years) in female murine liver TEM images. (M) Combined cristae score by sex across aging. (N-Q) Work through of 3D reconstruction by SBF-SEM. (N) Schematic depicting removal of the liver from mice. (O) Schematic showing Serial Block Face-Scanning Electron Microscopy (SBF-SEM) for serial imaging of ortho slices representing different depths of tissue. (P) Manual contour segmentation of ortho slices was performed to yield (Q) 3-dimensional (3-D) reconstructions of individually colored mitochondria. (R-S) Representative ortho slice images from (R) 3-month murine liver tissue and (S) 2-year murine liver tissue. (R’-S’) Representative ortho slice images with 3D reconstructions of mitochondria overlaid from (R’) 3-month murine liver tissue and (S’) 2-year murine liver tissue. (R’’-S’’) Isolated 3D reconstructions of mitochondria from (R’’) 3-month murine liver tissue and (S’’) 2-year murine liver tissue (T-Z) Quantification of 3-D reconstructed mitochondria for (T) mitochondrial perimeter, (U) area, and (V) volume in 3-month and 2-year-old mice (W-X) 3D reconstructions of mitochondria displayed from the transverse viewpoint in (W) 3-month and (X) 2-year murine liver tissue. (W’-X’) Representative images of 3D reconstructions of mitochondria displayed from the longitudinal viewpoint in (W’) 3-month and (X’) 2-year murine liver tissue. (Y-EE) Measurement of average mitochondrial sphericity (Y) and complexity index (Z) in 3-month and 2-year-old murine liver samples. AA-EE) Measurements of mitochondrial (AA) perimeter, (BB) 3D area, (CC) volume, (DD) sphericity, and (EE) complexity index in individual mice for 3-month (blue) and 2-year (pink) mice. Display of individual mitochondria as displayed by mito-otyping for (FF) 3-month and (GG) 2-year murine liver tissue. For SBF-SEM, a total of approximately 1500 mitochondria were analyzed, 750 per age group, from 3 mice per group. For all panels, error bars indicate SEM, Mann–Whitney tests were used for statistical analysis, and significance values are denoted as follows: **P ≤ 0.01, ****P ≤ 0.001, and ns, not significant.

With the 2D mitochondrial differences observed, we utilized 3D reconstruction of SBF-SEM images to further investigate changes in mitochondrial structure. For each age cohort, we analyzed approximately 250 mitochondria from each of the three mice surveyed (Figure 7O), totaling around 750 mitochondria per age point. SBF-SEM allowed for a resolution of 10 nm in the x- and y- axes, with 50 nm m intervals in the z-axis, to image a total of 300 slices. Of these, 50 slices were used for analysis. These slices underwent manual contour segmentation (Figure 7P) for the 3D reconstruction of mitochondria (Figure 7Q). Displayed first in both age cohorts are representatives of each orthoslice (Figure 7R and S), which allows for the identification of mitochondria (Figures 7R’ and S’). From there, we manually segmented each ortho slice in Amira to reconstruct mitochondria. Additionally, for better visualization of each individually colored mitochondria, the ortho slice may be removed (Figures 7R’’ and S’’). Once modeled, the dimensions of the 3D reconstructions can be quantified accurately. We found that in comparing the young and aged mice, mitochondrial size decreased significantly in all metrics (Figures 7T-X). Specifically, the perimeter, which represents the sum of all external distances in the mitochondria, was approximately 20% lower in aged samples (Figure 7T; 3 months: 10,761 ± 1,560 µm SD; 2 years: 8,054 ± 450 µm SD). Mitochondrial surface area, a metric of the outer mitochondrial membrane (OMM) area, was decreased by nearly 50% in aged samples (Figure 7U; 3 months: 7.66 ± 0.808 µm² SD; 2 years: 4.76 ± 0.511 µm² SD). Finally, mitochondrial volume, which represents the total of all internal pixels within the 3D mitochondrial reconstruction, was approximately 30% lower in aged male murine liver samples (Figure 7V; 3 months: 1.01 ± 0.182 µm³ SD; 2 years: 0.696 ± 0.107 µm³ SD). Losses in mitochondrial volume may indicate a decrease in internal volume for ATP synthesis,^75^ but mitochondrial roles extend beyond their energetics, including their ability to interface with the endoplasmic reticulum (ER) to modulate calcium homeostasis.^76^ Thus, it is equally important to look at their morphology and capacity to form contact sites with the ER.

To aid in visualizing mitochondria, we present mitochondria 3D reconstructions from transverse (Figures 7W and X) and longitudinal (Figures 7W’ and X’) viewpoints. To further verify this change across aging, we examined sphericity and the mitochondrial complexity index (MCI),^77^ which are analogous measures of mitochondrial morphology.

Sphericity (calculated as 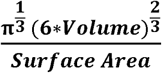), exhibited an approximate 20% higher mean value (Figure 7Y; 3 months: 0.638 ± 0.0211 SD; 2 years: 0.785 ± 0.0257 SD). MCI (calculated as Surface Area^3^ /16 π ^2^volume^2^) conversely showed a decrease in the aged cohort (Figure 7Z; 3 months: 3.22 ± 0.240 SD; 2 years: 1.64 ± 0.236 SD). Together, these findings validate decreased mitochondrial complexity in aged murine liver samples, with mitochondria adopting a more spherical morphology. To better visualize these changes, we organized mitochondria by their volume using a method known as mito-typing, which allows for comparison across different sizes (Figure FF and GG). In the three-month age cohort (Figure 5FF), we observed the typical morphology of mitochondria, characterized by a compact or tubular structure. However, we also observed some diversity in mitochondrial structure, such as elongation, branching, and other structures that prioritize surface area over volume. In contrast, the 2-year sample showed much less heterogeneity and mitochondrial structures were mostly presented as tubular or compact (Figure 7GG). Three-dimensional SBF-SEM analysis revealed significant age-associated changes in hepatic mitochondrial morphology. Compared with mitochondria from 3-month-old mice, those from 2-year-old mice exhibited reduced mitochondrial perimeter (7AA), area (7BB), volume (7CC), increased sphericity (7DD), and a decreased mitochondrial complexity index (Figure 7EE). Mito-otyping analysis further revealed that mitochondria from young liver tissue exhibited more elongated and morphologically diverse structures, whereas those from aged liver tissue appeared more spherical and less complex (Figure 7FF-GG). We based these analyses on approximately 1,500 mitochondria from three mice per age group. For all these metrics, while intra-sample heterogeneity is prevalent, there is minimal inter-sample heterogeneity and no outliers.

### Dietary and Metabolic Interplay with MICOS

Beyond aging, we wanted to further investigate the impact of a high-fat diet (HFD) on mitochondrial ultrastructure. We performed TEM analysis in a 20-week-old murine cohort subjected to a low-fat diet (LFD) and an HFD (Figures 8A-B’). Qualitatively, examination of TEM images revealed more lipid droplets with less circularity in HFD samples. Looking at mitochondria, although mitochondrial count, when normalized to the µm cell area, was lower in HFD samples (Figure 8C; LFD: 0.210 ± 0.114 SD; HFD: 0.128 ± 0.0972 SD; not significant). Similar to aging, HFD samples exhibit a significantly higher average mitochondrion area (Figure 8E; LFD: 0.570 ± 0.401 µm² SD; HFD: 0.923 ± 1.12 µm² SD). When considering these two quantifications in tandem, we also calculated the percentage of mitochondrial area related to the total cell area, which showed no significant difference (Figure 8D; LFD: 11.8 ± 5.59 µm² SD; HFD: 11.7 ± 5.10 µm² SD). When examining the changes in mitochondrial shape with an HFD, we observed that mitochondria had a higher circularity, indicative of potentially less complexity, compared to those with an LFD (Figure 8F; LFD: 0.781 ± 0.149 SD; HFD: 0.897 ± 0.0748 SD). Finally, examining the cristae score, similar to aging samples, HFD cohorts had a significantly lower cristae score (Figure 8G; LFD: 3.74 ± 0.441 SD; HFD: 2.04 ± 0.770 SD). Together, these findings show that an HFD can parallel age-related changes, and that an HFD exhibit smaller mitochondria with aberrant cristae. In addition, HFD induced significant metabolic alterations, including increased serum cholesterol (Figure 8J), serum triglycerides (Figure 8K), serum free fatty acids (Figure 8L), fat mass (Figure 8M), liver weight (Figure 8N), body weight (Figure 8O), serum insulin (Figure 8P) and OGTT values (Figure 8Q).

**Figure 8:**
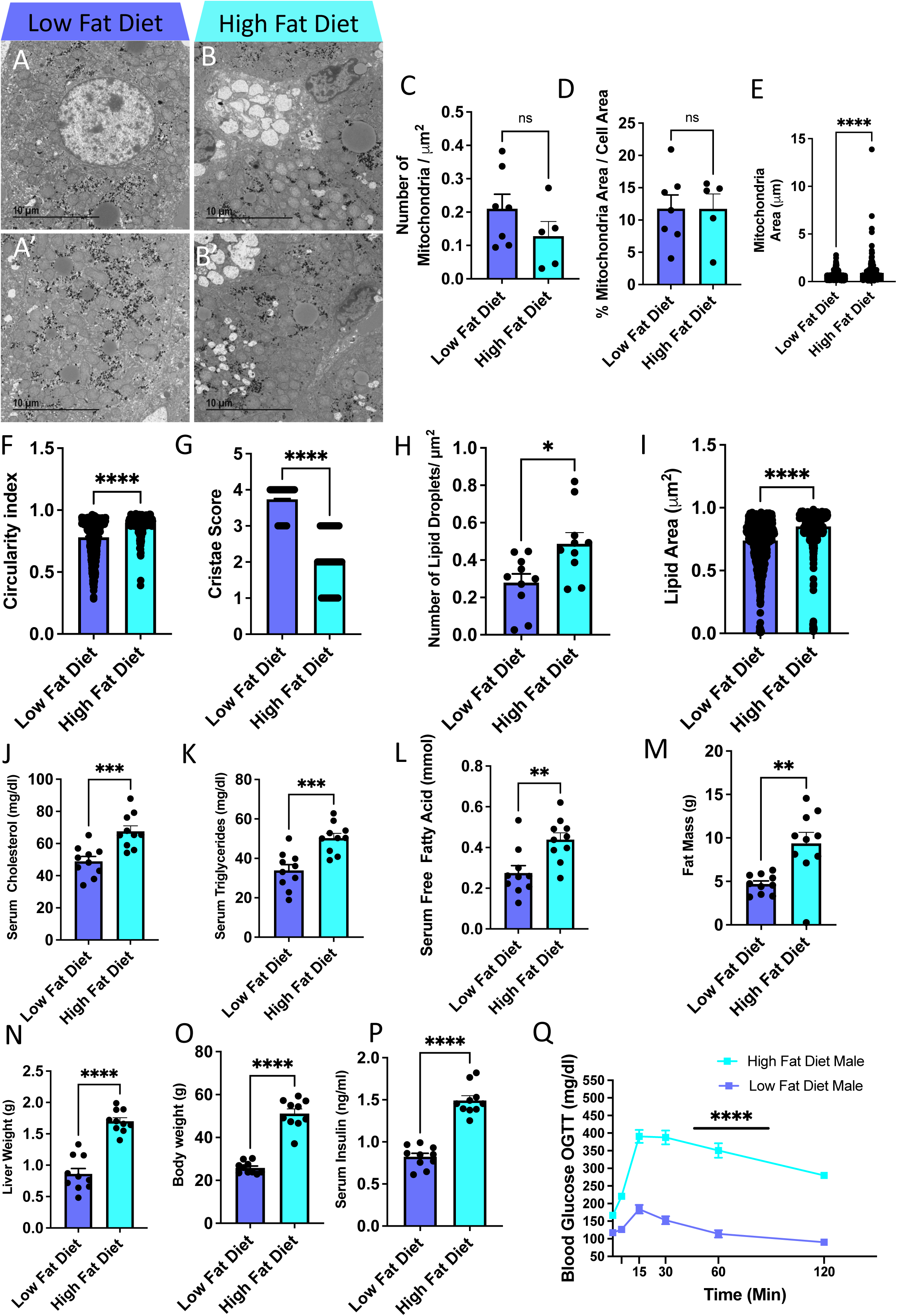
High-Fat vs. Low-Fat Diet Associated with Changes to Mitochondrial Morphology and Alterations to Physiological Parameters (A-B’) Representative transmission electron images of murine livers of mice fed (A-A’) low-fat and (B-B’) high-fat diets. (C-G) Measurements of (C) number of mitochondria per μm^2^ area, (D) percentage of mitochondrial area normalized to total cell area, (E) total mitochondrial area, (F) circularity index, and (G) cristae score of mitochondria from low-fat and high-fat diet murine livers. (H-I) Measurement of (H) number of lipid droplets per μm^2^ area and (I) lipid droplet area in low-fat and high-fat diet murine livers. (J-P) Measurement of physiological parameters (J) serum cholesterol, (K) serum triglycerides, (L) serum free fatty acids, (M) total fat mass, (N) liver weight, (O) body weight, and (P) serum insulin in mice fed low-fat and high-fat diets. (Q) Oral glucose tolerance test (OGTT) for male mice fed low-fat and high-fat diets. For all panels, error bars indicate SEM, ** indicates p< 0.01; and *p< 0.05, calculated with Student’s t-test.

## Discussion

### Genetic and Clinical Associations of MICOS Complex Genes with Liver Disease

First, we wanted to demonstrate a link between liver diseases and MICOS genes. Our study identified significant associations between genetically regulated expression (GREX) of MICOS complex genes and liver disease phenotypes within a biobank population. We observed that decreased GREX of *CHCHD3* (*MIC19*) in individuals of European ancestry correlated with liver cirrhosis and transplant status. In contrast, reduced *Opa1* expression in individuals of African ancestry was linked to chronic liver disease and cirrhosis. These findings highlight ancestry-specific genetic susceptibility to hepatic pathologies, potentially reflecting differences in genetic architecture or environmental interactions. However, the limitations of EHR-based phenotyping, such as incomplete data and diagnostic variability, underscore the need for replication in diverse cohorts. Future studies integrating longitudinal data and multi-omics approaches could elucidate how MICOS dysregulation interacts with aging and environmental factors to drive disease progression.

Next, we demonstrated that aging is associated with reduced expression of specific mitochondrial genes (*MIC19, MIC25,* and *MIC60*) and increased expression of *BIP, XBP1, IRE1,* and *ATF4*. The reduced expression of *MIC19, MIC25,* and *MIC60* has implications for mitochondrial structure and aging.^54,60^ As core components of the MICOS, *MIC19, MIC25*, and *MIC60* play a crucial role in maintaining cristae architecture, regulating oxidative phosphorylation, mitochondrial dynamics and bioenergetic function. ^22,24,78^ Reduced mRNA expression in older individuals is associated with loss of cristae integrity, which consequently reduces the surface area of respiratory complexes, thereby reducing mitochondrial efficiency & ATP production, and increasing susceptibility to oxidative damage from ROS.^16,54,60,109^ Thus, reduced *MIC19*, *MIC25* and *MIC60* likely reflects mitochondrial structural decay and reduced bioenergetic capacity with age.^114,115^

In aging cells, the upregulation of *BIP, IRE1, XBP1, ATF4,* and *CHOP* reflects a persistent activation of the endoplasmic reticulum (ER) stress response and the integrated stress response (ISR) as part of attempts to maintain cellular proteostasis under increasing proteotoxic and metabolic stress.^96,103^ *BIP* (also known as *GRP78*) acts as a key ER chaperone that dissociates from stress sensors upon accumulation of misfolded proteins, initiating the unfolded protein response (UPR) and signaling adaptive stress pathways.^96^ *IRE1* activation promotes the unconventional splicing of *XBP1* mRNA, resulting in the production of a potent transcription factor that enhances the expression of chaperones and ER quality control machinery.^103^ In contrast, *ATF4* is induced downstream of PERK signaling to regulate stress-responsive genes.^96^ Under prolonged or unresolved stress, *CHOP* expression increases, shifting the UPR from an adaptive to a pro-apoptotic signaling pathway by downregulating anti-apoptotic factors and linking to mitochondrial pathways of cell death.^96,103^ Collectively, elevated expression of these factors in aged tissues indicates chronic ER stress and an overwhelmed proteostasis network, which can disrupt ER–mitochondrial communication, exacerbate mitochondrial dysfunction, and contribute to age-related decline and susceptibility to disease.^82,83^

To validate and assess the importance of the MICOS genes in age-related changes, we knocked down *MIC60* and *CHCHD6*. We found that their deficiency impaired Ca^2+^ homeostasis, cell survival and induced oxidative stress in HepG2 cells.

### MICOS Complex Dysregulation Drives Mitochondrial Dysfunction

Previous studies have established that the MICOS complex is critical for mitochondrial dynamics,^78^ and our group have previously investigated the effect of aging on the MICOS complex in kidney tubular cells.^60^ As reported in our study, aging in murine liver was associated with a significant decline in the mRNA expression of key MICOS components and the associated protein OPA1. While *Opa1* interacts with the MICOS complex, it does not require *Opa1* to form cristae junctions, and loss of Opa*1* does not negatively affect MICOS components.^84^ This suggests that the loss of the MICOS complex across aging occurs in an *Opa1*-independent manner. This parallels our previous findings in skeletal muscle, the heart, and the kidney.^54,60,80^

Calcium influx in mitochondria can reflect cell viability.^97,98^ Mitochondrial mediation of calcium can impact various cell processes, including apoptosis, signaling, and ATP production.^99^ In some cases, elevated mitochondrial calcium uptake can occur antecedent to mitochondrial swelling, which results in a pathway often leading to apoptosis.^100^ Aging-associated declines in MICOS components (*MIC60, CHCHD3, CHCHD6*) correlated with disrupted calcium handling and elevated oxidative stress in HepG2 cells. Knockdown of *MIC60* or *CHCHD6* impaired mitochondrial calcium retention and increased ROS production, suggesting that MICOS integrity is vital for redox homeostasis and apoptosis regulation.^99,101^ This provides a plausible disease link as ER calcium release promotes mitochondrial dysfunction, inducing oxidative stress and hepatotoxicity,^102^ with broader implications in MASLD.^103,104^ Since Miro clusters both interact with the MICOS complex and regulate MERCs,^105^ Miro represents a potential future mechanistic avenue through which the MICOS complex contributes to MERC tethering.

Researchers have proposed that the mitochondrial redox state is a principal moderator of mitochondrial function in liver disease.^106^ Recent studies have continued to highlight a link between ROS and mitochondrial dynamics.^107^ Lipids can aid in stimulating the production of ROS.^108^ Additionally, many theories of aging propose that mitochondria lose function because ROS byproducts accumulate during aging.^109^ There remains controversy about whether ROS are generated in the liver. While some studies report that superoxides arise during liver aging,^110^ other studies conversely report that the aging human liver does not produce superoxides.^111^ Future studies should explore how ROS contributes to the observed 3D mitochondrial phenotypes, as oxidative stress may be a regulator of the phenotypes we observed. It is noteworthy that the loss of *CHCHD6* and *MIC60* leads to oxidative stress. This suggests that during aging, the liver undergo a vicious cycle wherein abnormal mitochondrial structures generate more harmful byproducts. These byproducts, in turn, worsen mitochondrial structural dysfunction, contributing to age-related oxidative stress. These findings align with studies linking cristae disorganization to mPTP sensitization and ATP synthase dysfunction.^102,103^ The vicious cycle of MICOS loss, oxidative stress, and metabolic dysfunction may accelerate age-related hepatic decline, particularly under lipid-rich conditions that exacerbate ER-mitochondrial uncoupling.^114,115^

### Hepatic Metabolic Reprogramming as a Feature of Aging

Our integrated metabolomic and lipidomic profiling of aged liver tissue reveals a profound and coordinated rewiring of hepatic metabolism, centering on mitochondrial dysfunction as a key driver of age-related physiological decline. The clear separation of young and aged metabolic signatures, along with the accumulation of vitamin A metabolites, suggests altered nutrient handling and redox signaling in aging hepatocytes.^17,18,62^ Concomitant reductions in tricarboxylic acid cycle intermediates, including succinate and malate, indicate impaired mitochondrial oxidative metabolism and/or increased cataplerotic flux, reflecting diminished bioenergetic capacity with age.^3,5,109,110^ In parallel, significant depletion of nucleotide monophosphates and NAD⁺-related metabolites highlights a decline in mitochondrial-dependent biosynthetic pathways and redox homeostasis, processes essential for energy production, cellular repair, and metabolic flexibility.^49,63,64^ Given the central role of mitochondria in coordinating these pathways, the observed metabolite changes collectively indicate compromised mitochondrial function as a key driver of altered energy metabolism, reduced anabolic capacity, and disrupted cellular homeostasis in the aging liver, thereby reinforcing mitochondrial dysfunction as a fundamental contributor to age-associated metabolic decline.^8,115^

Building upon the profound metabolic dysregulation observed, our findings demonstrate that age-related mitochondrial impairment in the liver has direct pathological consequences, culminating in overt lipid accumulation and a progression towards metabolic dysfunction-associated steatotic liver disease (MASLD).^20,66,116^ This transition is evident in both humans and mice, where advanced age is correlated with a significantly increased hepatic lipid content, exceeding established steatosis thresholds in human cohorts.^17,20,42^ In murine models, this manifests as a dramatic increase in lipid droplets and triglyceride concentration, accompanied by hallmark features of disease progression, such as a reduced liver-to-body weight ratio and decreased mtDNA content.^44,66,124^ Crucially, lipidomic profiling reveals that aging is not characterized by a uniform increase in lipids, but rather by a specific remodeling of the hepatic lipid landscape, including significant alterations in triglycerides, ceramides, and acylcarnitines.^68,69,70,71^ This specific signature links the compromised energy metabolism and NAD^+^ depletion which impair fatty acid oxidation, to a lipogenic shift and the accumulation of lipid species known to promote insulin resistance and cellular stress.^66,69,102,108^ Thus, age-related mitochondrial dysfunction appears to be a central driver that shifts the liver from a state of metabolic flexibility to one of lipid-laden inflexibility, significantly increasing the risk of serious hepatic and systemic metabolic disease.^114,115,116^

### Age-Related Mitochondrial Structural Remodeling in the Liver

Using SBF-SEM, we demonstrated that aging reduces mitochondrial volume, perimeter, and complexity in the murine liver, paralleling 2D TEM findings of cristae disorganization and fragmentation. These structural declines are associated with impaired metabolic and lipidomic profiles, indicating compromised bioenergetic efficiency and oxidative capacity. A limitation of our study is the lack of direct measurement of ATP synthesis rates, which would have more directly linked structure to function; this is an important area for future investigation. Notably, aged mitochondria exhibited spherical morphologies with reduced contact site potential, which may disrupt mitochondria-ER interactions critical for lipid homeostasis and calcium signaling.^82–84^ The structure of mitochondria remains relevant in aging as structural decline may decrease mitochondrial function, thereby impairing certain liver roles. While megamitochondria, a hallmark of advanced liver disease,^85–87^ were absent in our aged cohort, this may reflect a pre-diseased state where compensatory mechanisms, such as upregulated *Drp1*, remain functional.^86,88,89^ The observed structural changes likely precede overt pathology, underscoring the role of mitochondrial integrity in maintaining hepatic resilience during aging.

Beyond mitochondrial 3D structure, past studies have shown that fatty acid accumulation promote endoplasmic reticulum stress and liver injury in rodent models.^90^ Due to the increase of lipid droplets in both aging and high fat diet treatment in TEM images, there may be an increase in mitochondria-lipid droplet contact sites (MLDCs).^76,91^ Beyond this, lipid droplet-endoplasmic reticulum contacts are also understood to help form lipid droplets and perform metabolism.^65^ Especially relevant is that MLDCs serve as a place for fatty acid homeostasis.^91^ Additionally, mitochondria endoplasmic reticulum contact sites (MERCs) importantly play a role in lipid homeostasis and synthesis.^92^ MERCs are understood to regulate ER stress,^93^ so increases in fatty acids, triggered by lipid droplet formation, can lead to increased MERC formation in the aged model. Conversely, we qualitatively observed a decrease in structures consistent with wrappER, a mitochondria-ER contact known to maintain lipid flux through a tri-organelle contact site that involves peroxisomes.^94^ These wrappER sites, which contain sites of adhesion, regulate very-low-density lipoproteins.^95^ The age-related loss of mitochondrial complexity may impair the ability and relative surface area mitochondria in murine liver samples must form contact sites, including MERCs, thus interfering with functions including lipid homeostasis. Indeed, past reviews have suggested targeting MERCs in MASLD due to the role of contact sites in glucose and lipid metabolism.^96^ However, in the future, a more rigorous analysis of age-related changes in contact sites within liver tissue is necessary.

### Dietary and Metabolic Interplay with MICOS in Aging

Our investigation into the impact of a high-fat diet reveals that nutritional stress recapitulates and accelerates the same fundamental mitochondrial pathology observed in aging, directly linking dietary patterns to premature cellular aging and metabolic dysfunction. The HFD-induced mitochondrial remodeling characterized by enlarged yet simplified organelles with significantly disordered cristae architecture mirrors the ultrastructural decay seen in aged livers, suggesting a common endpoint of bioenergetic compromise. This loss of cristae integrity is particularly critical, as it directly impairs the capacity for oxidative phosphorylation. While the total mitochondrial area per cell was maintained, the shift towards fewer, larger, and more circular organelles points to an imbalance in mitochondrial dynamics, likely hindering their ability to network, traffic, and undergo efficient quality control. These structural deficits manifest in severe systemic metabolic alterations, including elevated serum lipids, insulin resistance, and hepatic steatosis. Therefore, HFD does not merely alter metabolism passively but actively drives a pathological transformation of mitochondrial structure that underpins the development of metabolic disease, positioning poor dietary quality as a potent accelerator of the mitochondrial dysfunction typically associated with advancing age.

We know that diet induced obesity (DIO) are linked with many liver complications. For example, oxidative stress, inflammation, and lipogenesis are factors that may be linked to DIO and exacerbate the onset of MASLD.^116^ Additionally, DIO is associated with metabolic dysregulation of the liver.^117^ Elevated retinol and ceramide levels in aged liver paralleled HFD-associated lipotoxicity, implicating disrupted β-oxidation, impaired insulin sensitivity and peroxisomal signaling.^17,118^ The enhanced supply of vitamin A and high-fat consumption in diet-induced obese mice is linked to the production of bisretinoid.^119^ Interestingly, obesity within hepatocytes has also been shown to increase MAMs (i.e., MERC-isolated biochemical fractions), which confers mitochondrial dysfunction.^120,121^ Thus, HFDs can play a role in metabolic defects in the liver tissue, although this requires further investigation in the future.

An HFD can lead to pathological alterations and damage to the ultrastructure of the mitochondria, as well as downregulation of MERCs regulators, *MFN2* and *OPA1*^122^ which parallels the TEM ultrastructural changes we observed. Additionally, studies have shown that *Mic19/Chchd3* is required for normal liver function. In the liver, deletion of *Mic19/Chchd3* reduces ER-mitochondrial contacts, disrupts mitochondrial lipid metabolism, disorganizes mitochondrial cristae, and triggers an unfolded protein stress response in mouse hepatocytes, resulting in impairments of liver mitochondrial fatty acid β-oxidation and lipid metabolism.^123^ Thus, while MICOS components like *MIC19* and *CHCHD3* are essential for ER-mitochondrial tethering and lipid metabolism; their age-related decline may impair compensatory responses to dietary stress. Notably, HFD and aging both reduce mitochondrial plasticity, exacerbating steatosis and insulin resistance, key drivers of MASLD progression.^124,125^

## Conclusions

Our integrative analysis reveals that age-related disruption of the MICOS complex contributes to mitochondrial structural and functional decline in the liver, promoting metabolic dysfunction and increasing disease susceptibility. By linking 3D ultrastructural changes, genetic associations, and cellular mechanisms, we establish MICOS as a critical node in the process of hepatic aging. A limitation of this study is the primary use of a male murine model for deep ultrastructural and functional validation; future work should explore sex-specific differences in these processes. Deleting the MICOS complex in HepG2 cells impairs calcium uptake and increases oxidative stress, underscoring the importance of this complex. A high-fat diet exacerbates lipid synthesis, catabolism, and mitochondrial changes in aging livers, increasing susceptibility to liver disease.^17,116^ This study demonstrates that impaired mitochondrial structure, calcium influx, ROS production, metabolic alterations, and MICOS dysfunctions contribute to liver diseases. These findings highlight potential therapeutic targets, such as enhancing cristae integrity or modulating MERCs, to mitigate age-related liver pathologies. Future studies examining tissue-specific MICOS dynamics across diverse populations and dietary conditions will further elucidate its role in aging and metabolic diseases.

## Supporting information

CTAT methods

## Abbreviations

TEM, transmission electron microscopy; SBF-SEM, serial block-face scanning electron microscopy; MASLD, metabolic dysfunction-associated steatotic liver disease; MICOS, mitochondria contact site and cristae organizing system; GREX, genetically regulated gene expression; MRI, magnetic resonance imaging; TCA, tricarboxylic acid; NAD^+^, nicotinamide adenine nucleotide; TGO, triglyceride oligomer; TG, triglycerides; Cer, ceramide; CAR, acylcarnitine; ER, endoplasmic reticulum; MCI, mitochondrial complexity index; ROS, reactive oxygen species; MLDC, mitochondria-lipid contact site; HFD, high-fat diet; DIO, diet induced obesity; MAM, mitochondria-associated membrane; MERC, mitochondria endoplasmic reticulum contact site; OMM, Outer Mitochondrial Membrane.

## Authors’ contributions

Sepiso K. Masenga, Alexandria Murphy, Prasanna Venkhatesh contributed equally to this work. Zer Vue, Benjamin Rodriguez, Ashlesha Kadam, Andrea G. Marshall, Estevão Scudese, Brenita Jenkins, Amber Crabtree, Praveena Prasad, Edgar Garza-Lopez, Han Le, Ky’Era V. Actkins, Elma Zaganjor, Nelson Wandira, Jeremiah Afolabi, Prasanna Katti, Chantell Evans, Young Do Koo, Dhanendra Tomar, Mark A. Phillips, David Hubert, Chandravanu Dash, Pooja Jadiya, Olujimi A. Ajijola, Magdalene Ameka, Okwute M. Ochayi, Eric Wang, Quinton Smith, Ronald McMillan, Annet Kirabo, André Kinder, Tyne W. Miller-Fleming, Bret Mobley, Julia D. Berry, Nathan Winn, Vernat Exil, Anita M. Quintana, Kit Neikirk, Jenny Schafer, Sean Schaffer, Oleg Kovtun, Mohd Mabood Khan, Calixto Pablo Hernandez Perez, Margaret Mungai, Melanie R. McReynolds, Antentor Hinton, Jr contributed to the conception, design, data acquisition, analysis, and interpretation of data. Melanie R. McReynolds and Antentor Hinton, Jr. conceived and supervised this study. All authors were involved in drafting and critically revising the manuscript for important intellectual content. All authors approved the final version of the manuscript.

## Data availability statement

All data generated or analyzed during this study are included in this published article and its Supplementary information files. Additional data can be requested from the corresponding author.

## Acknowledgment

We would like to acknowledge the Huck Institutes’ Metabolomics Core Facility (RRID:SCR_023864) for use of the OE 240 LCMS and Sergei Koshkin for helpful discussions on sample preparation and analysis. We would also like to acknowledge the Huck Institutes’ Metabolomics Core Facility (RRID:SCR_023864) for use of the OE 240 LCMS and Drs. Imhoi Koo, Ashley Shay, and Sergei Koshkin for helpful discussions on sample preparation and analysis. We would also like to thank UCLA investigators for gifting us old and young human liver samples.

