## Supplementary material for "Age-related MICOS Complex Dysregulation Impairs Mitochondrial 3D Architecture and Metabolic Homeostasis in the Liver": CTAT methods

Tables for a “Complete, Transparent, Accurate and Timely account” (CTAT) are now mandatory for all revised submissions. The aim is to enhance the reproducibility of methods.

- Only include the parts relevant to your study
- Refer to the CTAT in the main text as ‘Supplementary CTAT Table’
- Do not add subheadings
- Add as many rows as needed to include all information
- Only include one item per row

**If the CTAT form is not relevant to your study, please outline the reasons why:**

1.1 **Antibodies**

| **Name** | **Citation** | **Supplier** | **Cat no.** | **Clone no.** |
| --- | --- | --- | --- | --- |
| **Anti-Mitofilin antibody [2E4AD5] - Mitochondrial Marker** | **Validated in Tilokani, L., Russell, F. M., Hamilton, S., Virga, D. M., Segawa, M., Paupe, V., Gruszczyk, A. V., Protasoni, M., Tabara, L. C., Johnson, M., Anand, H., Murphy, M. P., Hardie, D. G., Polleux, F., & Prudent, J. (2022). AMPK-dependent phosphorylation of MTFR1L regulates mitochondrial morphology. Science advances, 8(45), eabo7956. https://doi.org/10.1126/sciadv.abo7956** | **abcam** | **ab110329** | 2E4AD5 |
| **Sam50 Polyclonal antibody** | **Validated in Guarani, V., McNeill, E. M., Paulo, J. A., Huttlin, E. L., Fröhlich, F., Gygi, S. P., Van Vactor, D., & Harper, J. W. (2015). QIL1 is a novel mitochondrial protein required for MICOS complex stability and cristae morphology. eLife, 4, e06265. https://doi.org/10.7554/eLife.06265** | **proteintech** | **20824-1-AP** |  |
| **alpha Tubulin Antibody (DM1A) - BSA Free** | **Validated in** [Fazel M, Wester MJ, Schodt DJ et al. High-precision estimation of emitter positions using Bayesian grouping of localizations Nature communications 2022-11-22 [PMID: 36418347]](http://www.ncbi.nlm.nih.gov/pubmed/36418347) | **Novus** | NB100-690 | DM1A |
| **Donkey anti-Mouse IgG (H+L) Highly Cross-Adsorbed Secondary Antibody, Alexa Fluor™ Plus 800** | **Validated in** 3D reconstruction of murine mitochondria reveals changes in structure during aging linked to the MICOS complex. | **Invitrogen** | A32789 |  |
| **Donkey anti-Rabbit IgG (H+L) Highly Cross-Adsorbed Secondary Antibody, Alexa Fluor™ Plus 680** | **Validated in Validating Enteroid-Derived Monolayers from Murine Gut Organoids for Toxicological Testing of Inorganic Particles: Proof-of-Concept with Food-Grade Titanium Dioxide.** | **Invitrogen** | A32802 |  |

1.2 **Cell lines**

| **Name** | **Citation** | **Supplier** | **Cat no.** | **Passage no.** | **Authentication test method** |
| --- | --- | --- | --- | --- | --- |
| *HepG2* | **Validated in Lieber A, et al. Recombinant adenoviruses with large deletions geneerated by cre-mediated excision exhibit different biological properties compared with first-generation vectors in vitro and in vivo. J. Virol. 70: 8944-8960, 1996. PubMed: 8971024** | **ATCC** | **HB-8065** | **3** | **Checking for proper morphology by microscopy** |

1.3 **Organisms**

| **Name** | **Citation** | **Supplier** | **Strain** | **Sex** | **Age** | **Overall n number** |
| --- | --- | --- | --- | --- | --- | --- |
| **C57BL/6** | **Validated in Expression of the Neuronal tRNA n-Tr20 Regulates Synaptic Transmission and Seizure Susceptibility.** | **The Jackson Laboratory in Maine** | **000664** | **Male** | **3-month, 2 year** | **6** |

1.4 **Sequence based reagents**

| **Name** | **Sequence** | **Supplier** |
| --- | --- | --- |
| **Mic19 (Human)** | **Forward- 5’-GAAAAGAATCCAGGCCCTTCCACGCGC-3’**  **Reverse- 5’-CAGTGCCTAGCACTTGGCACAACCAGGAA-3’** | **IDT** |
| **Mic25 (Human)** | **Forward- 5’-CTCAGCATGGACCTGGTAGGCACTGGGC-3’**  **Reverse- 5’-GCCTCAATTCCCACATGGAGAAAGTGGC-3’** | **IDT** |
| **BIP (Human)** | **Forward- 5’-CTGTCCAGGCTGGTGTGCTCT-3'**  **Reverse- 5’-CTTGGTAGGCACCACTGTGTTC-3'** | **Origene** |
| **PERK (Human)** | **Forward- 5’-GTCCCAAGGCTTTGGAATCTGTC-3'**  **Reverse- 5’-CCTACCAAGACAGGAGTTCTGG-3'** | **Origene** |
| **XBP1 (Human)** | **Forward- 5’-CTGCCAGAGATCGAAAGAAGGC-3'**  **Reverse- 5’-CTCCTGGTTCTCAACTACAAGGC-3'** | **Origene** |
| **CHOP (Human)** | **Forward- 5’-GGTATGAGGACCTGCAAGAGGT-3'**  **Reverse- 5’-CTTGTGACCTCTGCTGGTTCTG-3'** | **Origene** |
| **Mic60 (Human)** | **Forward- 5’-CCTCCGGCAGTGTTCACCTAGTAACCCCTT-3’**  **Reverse- 5’-TCGCCCGTCGACCTTCAGCACTGAAAACCTAT-3’** | **IDT** |
| **IRE1 (Human)** | **Forward- 5’-CCGAACGTGATCCGCTACTTCT-3'**  **Reverse- 5’-CGCAAAGTCCTTCTGCTCCACA-3'** | **Origene** |
| **ATF4 (Human)** | **Forward- 5’-AACCTCATGGGTTCTCCAGCGA-3'**  **Reverse- 5’-CTCCAACATCCAATCTGTCCCG-3'** | **Origene** |
| **Opa1 (Mouse)** | **Forward- 5’-TCTCAGCCTTGCTGTGTCAGAC-3'**  **Reverse- 5’-TTCCGTCTCTAGGTTAAAGCGCG-3'** | **Origene** |
| **Mitofilin (Mouse)** | **Forward- 5’- CAGTTCAGGCTGTCAAGGCACA-3'**  **Reverse- 5’- CATCAACTGCCTTTCTTCGCTCC-3'** | **IDT** |
| **Chchd3 (Mouse)** | **Forward- 5’- GCGGACGAGAACGAGAACAT-3'**  **Reverse- 5’- TCGCTGAGACTTAGAGCCAGA-3'** | **IDT** |
| **Chchd6 (Mouse)** | **Forward- 5’- TGTCTGAAAGTGTTGTGAACCG-3'**  **Reverse- 5’- GATGGCTGGTAGAGGGACAGT -3'** | **IDT** |
| **Cox1 (Mouse)** | **Forward- 5’- GCCCCAGATATAGCATTCCC-3'**  **Reverse- 5’- GTTCATCCTGTTCCTGCTCC-3'** | **IDT** |
| **Rpl13a (Mouse)** | **Forward- 5’- GAGGCCCCTACCATTTCCGA-3'**  **Reverse- 5’- GGCTTCAGCCGAACAACCTT-3'** | **IDT** |

1.5 **Biological samples**

| **Description** | **Source** | **Identifier** |
| --- | --- | --- |
| **Human exercise data** | Autopsy cases from Brazilian cohorts | CAEE number 77570224.2.0000.5281 |

1.6 **Deposited data**

| **Name of repository** | **Identifier** | **Link** |
| --- | --- | --- |
| **GitHub** | **Lipidomic-Analysis-Young-and-Aged-Mouse-Tissue** | **https://github.com/mphillips67/Lipidomic-Analysis-Young-and-Aged-Mouse-Tissue** |

1.7 **Software**

| **Software name** | **Manufacturer** | **Version** |
| --- | --- | --- |
| **Amira** | **Thermofisher** | **2023** |
| **GraphPad Prism** | **GraphPad Software** | **10.4.0** |

1.8 **Other (e.g. drugs, proteins, vectors etc.)**

| Lipofectamine RNAiMax (Invitrogen) | MIC60 siRNA (Invitrogen) | CHCHD6 siRNA (Invitrogen) |
| --- | --- | --- |

1.9 **Please provide the details of the corresponding methods author for the manuscript:**

**2.0	Please confirm for randomised controlled trials all versions of the clinical protocol are included in the submission. These will be published online as supplementary information.**
